# Differential bioactivity of four BMP-family members as function of biomaterial stiffness

**DOI:** 10.1101/2021.02.10.430282

**Authors:** Adrià Sales, Valia Khodr, Paul Machillot, Laure Fourel, Amaris Guevara-Garcia, Elisa Migliorini, Corinne Albigès-Rizo, Catherine Picart

## Abstract

Whereas soft biomaterial is not able to induce cell spreading, BMP-2 presented by a soft film has been described to be sufficient to trigger cell spreading, migration and downstream BMP-2 signaling. Based on thin polyelectrolyte films of controlled stiffness, we investigated whether the presentation of four BMP members (2, 4, 7, 9) in a matrix-bound manner may differentially impact cell adhesion and bone differentiation of skeletal progenitors. We performed high content and automated screening of cellular responses, including cell number, cell spreading area, SMAD phosphorylation and alkaline phosphatase activity. The basolateral presentation of the different BMPs allowed us to discriminate the specificity of cellular response and the role of BMP receptors type I, type II, as well as three β integrins, in a BMP type and stiffness-dependent manner.

## Introduction

Bone morphogenetic proteins (BMPs) belong to the transforming growth factor β superfamily ^1^ and are known to participate in a large number of physiological and pathological processes, including cell growth and differentiation ^2^, tissue development ^3^, as well as cancerous processes ^4,5^. BMP signaling, which is mediated via BMP type I and type II receptors, involves different pathways usually described as SMAD or non-SMAD, and is dependent on the mechanical context of the cell environment ^6^. BMP signaling pathway is under the control of multiple ligands binding two type I and two type II receptors that interact combinatorically to form distinct receptor-ligand complexes ^7^. In addition, the repertoire of BMP ligands and of BMP receptors is cell- and tissue-dependent context ^8^.

BMPs can either be diffusible molecules or be presented by extracellular matrix (ECM) components ^9^. It is known that BMP-mediated signaling depends on the duration of exposure to BMPs and on their spatial localization ^10,11^. A sustained presentation of the BMP ligands can be obtained using biomaterials such as a collagen ^12^, a poly(ethyl acrylate) polymer ^13^, or a polyelectrolyte film containing hyaluronic acid and poly(L-lysine) ^14^.

Beside their role in cell differentiation, BMPs appear to induce cell spreading and they also appear to control cytoskeleton organization and cell migration ^15,16^. Increasing evidences highlight a role of BMPs in mechanotransduction ^6,17,18^. Mechanotransduction via the ECM is known to involve the adhesion receptors integrins and to be sensitive to matrix stiffness ^19,20^. Indeed, BMP signaling can be modulated by integrins ^21^, matrix stiffness and cytoskeletal tension ^22^.

We have previously shown that a biomimetic material presenting BMP-2 in a matrix-bound manner (bBMP-2) can be used to control stem cell fate *in vitro* ^23,24^ and *in vivo* to repair critical-sized bone defects ^25^. We have also shown that bBMP-2 presented from a soft film was sufficient to trigger cell spreading and migration ^26^, overriding the response to film stiffness. The same effect has been shown on rigid surfaces with a low cRGD surface density ^27^ and on surfaces presenting peptides specific for α5β1 integrins ^28^. This process on soft films involved β3 integrins and BMP receptors that worked together to control SMAD signaling and tensional homeostasis^26^, thereby coupling cell adhesion and fate commitment. Whether other BMPs are able to induce BMP receptor-integrin crosstalk is poorly known. For instance, BMP-4, which is involved in hematopoiesis, leukemias and cancers ^29^, is known to induce the expression of α4-integrin ^16^. BMP-7 plays a major role in fibrosis, kidney disease and is involved in the adhesion and migration of human monocytic cells ^30^. BMP-9 is involved in cardiovascular diseases and anemia ^18^. It also increases cell proliferation on bone grafts ^31^, but its overexpression decreases cell adhesion and migration ^32^. However, whether the signaling induced by BMPs is stiffness-dependent, is still elusive.

Recently, we have developed an automated process to deposit the polyelectrolyte films in various multiwell plate formats, especially 96-well microplates ^24^. Interestingly, such film-coated microwells enable to study a large number of experimental conditions in parallel. In addition, multiwell plates are compatible with automated methods to quantify cellular processes using plate readers and automated imaging systems.

In this study, four BMPs including BMP-2, −4, −7 and −9 were selected in view of their physiological importance such as skeletal formation, involvement in cardiovascular diseases, cancers, hematopoiesis and leukemias ^2,5,18,29^. All four BMPs have an osteogenic potential *in vitro* ^33^. We varied film stiffness to assess the combined effect of bBMP type and substrate mechanics on cell adhesion and differentiation. As BMP-responsive cells, we selected C2C12 skeletal myoblasts ^33,34^ and human periosteum-derived stem cells (hPDSCs) ^35,36^ in view of their physiological relevance for bone regeneration and known sensitivity to a large number of BMPs, including these four BMPs ^33,37^.We directly compared the effect of the four selected BMPs, on stem cell adhesion and early bone differentiation in response to increasing doses of bBMPs presented by the polyelectrolyte film. Cell adhesion, assessed by the number of adherent cells and the cell spreading area, and early bone differentiation, assessed by quantifying SMAD phosphorylation and alkaline phosphatase (ALP) activation, were quantified in an automated manner. Finally, by means of silencing RNA against the type I and II BMP receptors and against three selected β integrins, we identify the roles of each BMP receptor and integrins in cell adhesion, SMAD signaling and ALP expression in BMP ligand- and film stiffness-dependent context.

## Results

### Cell adhesion and spreading of C2C12 and hPDSCs is induced in a BMP-specific and stiffness-dependent manner

To investigate whether the film stiffness may affect the effect induced by the four BMPs, namely BMP-2, −4, −7 and −9, in early cell adhesion and spreading we bound them non-covalently to a polyelectrolyte film of controlled stiffness made of poly(L-lysine) (PLL) and hyaluronic acid (HA) ^14,26,38^. The polyelectrolyte films were crosslinked to different levels using a crosslinker EDC, as previously described ^38^, which leads to low and high crosslinked films (EDC30 and EDC70, respectively), named hereafter for simplicity soft (S) and rigid (R) films. Their stiffness corresponds to ~200 kPa (soft) and ~400 kPa (rigid) ^38^. The amounts of the four BMPs loaded in the films were quantified (**Table SI 1**). BMP-2, −4, −7, and −9 could all be loaded in the films at 65 to 95% of the loading concentration depending on the BMP type, with only slight stiffness-differences. Surprisingly, BMP-9, which is known to be found circulating in plasma in soluble form ^39^, was well bound to the matrix, with incorporation percentages of 92 and 88% on soft and rigid films, respectively. Thus, the four BMPs could be presented to cells in a matrix-bound manner (bBMPs).

Cells were fixed and analyzed 4-5 h after cell seeding to enable cell adhesion and spreading on the soft and stiff films. Both cell processes are usually slower on the films than on tissue culture polystyrene or glass, both of which are orders of magnitude stiffer than the films ^23^. Soluble BMPs (sBMPs) in the cell culture medium were used as control, at a constant concentration of 200 ng/ml, which is considered as a high concentration ^23^. Cells were cultured on soft and rigid films with increasing concentrations of bBMPs, obtained by loading the films with BMP solutions from 0 to 20 μg/mL. Representative images of cell adhesion are given in **Fig. 1A** and **Fig. SI 1**, together with the quantification of the number of adherent cells (**Fig. 1B**) and cell spreading areas (**Fig. 1C**). At first sight, it appears that the cells adhere and spread on all bBMPs in a concentration-dependent and stiffness-dependent manner, whatever the bBMPs, except on soft films with bBMP-9. After quantitative analysis, we observe that cell number and cell area values are generally higher on rigid films than on soft ones (**Fig. 1B** and **C**). However, on the non-BMP condition, this increase is only observed for cell number but not for cell spreading.

**FIGURE 1.**
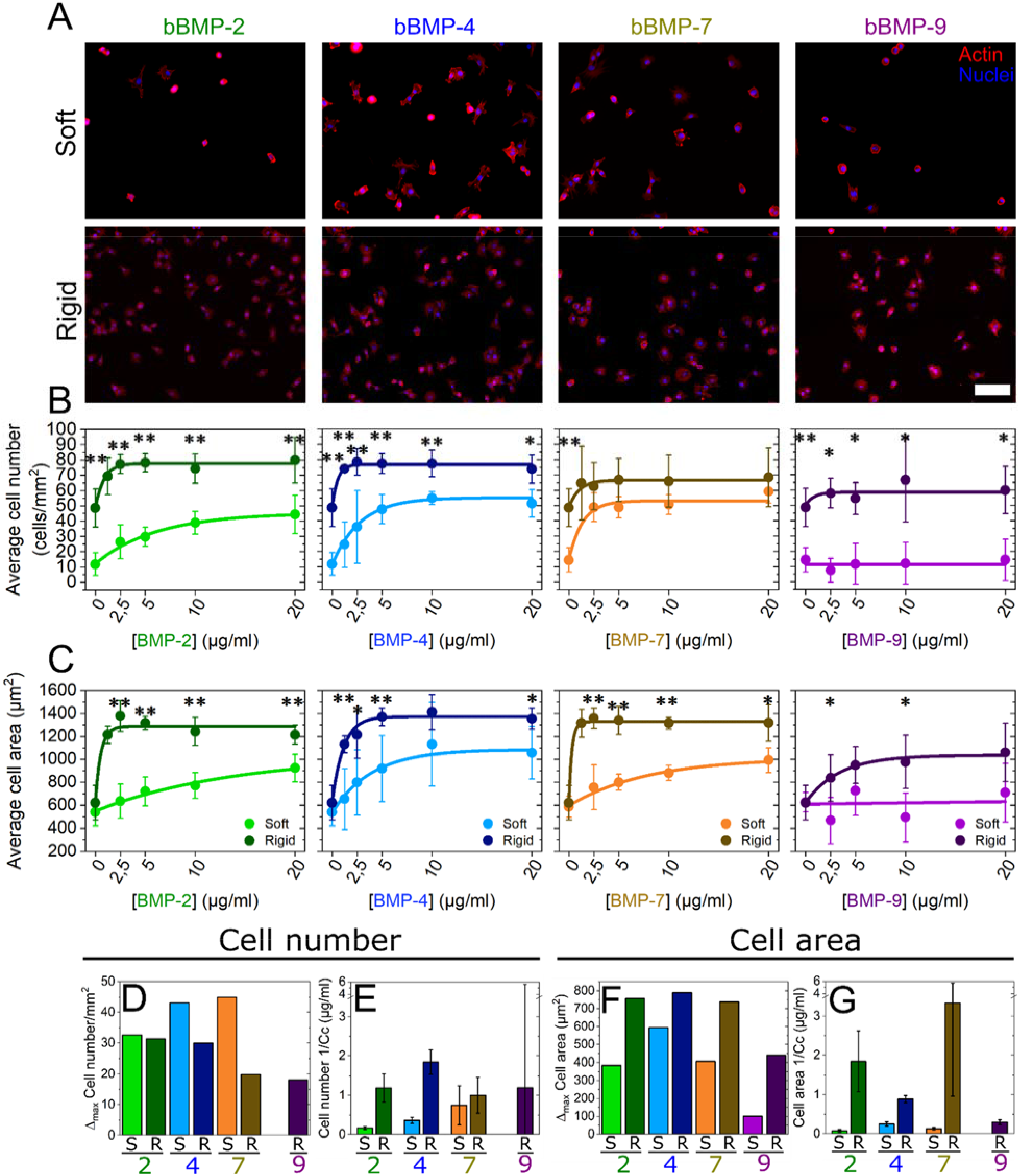
Quantification of C2C12 cell adhesion on soft and rigid films with four bBMPs (2, 4, 7, 9) at increasing BMP-loading concentrations. Representative fluorescent images of the actin cytoskeleton for (**a**) cells cultured for 5 h on soft films (S) and for 4 h on rigid films (R). Images correspond to a BMP loading concentration in solution of 5 μg/mL. Quantification of (**b**) the average cell number per mm^2^ of substrate area and **(c**) the cell area as a function of the initial BMP concentration in the loading solution. Results of the data fitting for each BMP-dose response curve (see methods) gave two parameters: Δ_max_ and Cc (characteristic concentration). (**d**) Difference (Δ_max_) between the maximum cell number and the no bBMP condition, plotted for each of the 4 bBMPs and film stiffness (S, R films). (**e**) Sensitivity to the BMP concentration (given as 1/Cc that provides a direct information on the sensitivity to the bBMP concentration). (**f** and **g**) Same quantitative parameters extracted for the cell spreading area in response to the bBMP-dose. Data represent three independent biological replicates with two microwells per condition in each independent experiment (technical replicates). Statistical tests were done using non-parametric Kruskal-Wallis ANOVA test to compare S (light color) and R (dark color) films (*p < 0.05; **p < 0.01). Scale bar=100 μm.

The quantified data were fitted with an exponential function toward a plateau value (see *Methods*) and two quantitative parameters were extracted: Δ_max_ (difference between the highest value and value for the no BMP condition) and 1/Cc (**Fig. 1D-G**). Cc (in μg/mL) corresponds to the characteristic concentration extracted from the exponential fit of the data. It corresponds approximately to the BMP concentration when half of the plateau value is reached. 1/Cc provides a direct quantification of the cell sensitivity to bBMPs: the higher it is, the higher is the sensitivity to a given bBMP. Regarding Δ_max_, the higher it is, the stronger is the effect for a given BMP.

The increase in cell number (Δ_max_) was stiffness-dependent for bBMP-4 and −7, being higher on soft than on rigid films (**Fig. 1D**). In cell area, this same parameter was found to be stiffness-dependent for all the bBMPs. However, the increase in cell area was always higher on rigid films than on soft ones (**Fig. 1F**). Regarding the sensitivity parameter (1/Cc), it was stiffness-sensitive for bBMP-2 and −4 in cell number (**Fig. 1E**), and for bBMP-2, −4 and −7 in cell area (**Fig. 1G**), being in all cases higher on rigid films than on soft ones. However, a rather large standard deviation was found for bBMP-7 sensitivity parameter on cell area. bBMP-9, in particular, only helped increasing adhesion and spreading on rigid films and no response was observed on soft ones at this time scale (**Fig. 1B** and **D**), indicating a smaller role on adhesion and spreading of this bBMP in comparison to the others.

Interestingly, plotting the data in relative percentage compared to the no BMP condition, further highlighted the fact that soft films with bBMPs had a stronger role on cell adhesion than rigid films, while rigid films with bBMPs had a stronger role on cell spreading (**Fig. SI 2**). Soluble BMPs (sBMPs) added on cells cultured in the same experimental conditions than with bBMPs, only had an effect increasing cell area on rigid films with sBMP-2 (**Fig. SI 3**), confirming that bBMPs amplify cell adhesive response.

The adhesive response of hPDSCs (**Fig. SI 4** and **SI 5)** was qualitatively and quantitatively similar to that of C2C12 myoblasts. However, we noted some differences: regarding the Δ_max_ value, cell adhesion was stiffness-dependent for all four BMPs (**Fig. SI 3**) while the cell area was only stiffness-dependent for bBMP-4 and higher on soft than on rigid films (**Fig. SI 3G**). Regarding the kinetics, faster adhesion kinetics were observed on rigid films with bBMP-7 in comparison to soft ones (**Fig. SI 3F** and **H**), while no stiffness-dependent differences were observed on C2C12 cells. We also noted that, on rigid films with bBMP-2, bBMP-7, and to a lesser extent with bBMP-4, the cell number and cell area values, decreased after reaching the maximum value, showing a dual response of hPDSCs to high bBMP concentrations. Finally, on bBMP-9, contrary to C2C12 cells, very small adhesion could be measured on soft films (**Fig. SI 3B**). As with C2C12, data in relative percentage further highlighted the fact that soft films with bBMPs had a strong role of both cell adhesion and spreading of hPDSCs (**Fig. SI 5**). Here again, this effect was specific to bBMPs since sBMPs only had an influence increasing cell area on rigid films with sBMP-2 (**Fig. SI 6**).

Altogether, our data showed that the four studied bBMPs positively act either on the cell number, on the spreading area or both. There is BMP-specific response to film stiffness. The increase in cell number thanks to bBMPs was stronger on soft films than on rigid ones, notably for BMP-4 and −7, while cell area was more strongly enhanced by bBMPs on rigid than soft films. In general, cells respond faster, in terms of adhesion and spreading, on rigid films than soft ones. Similar behaviors were observed between C2C12 cells and hPDSCs.

### Early stem cell differentiation in bone is BMP-specific and stiffness-dependent

SMAD1,5,9 is a transcription factor known to play a key role in the transduction pathway from BMP receptors to the nucleus ^40^ and its phosphorylation is often taken as readout of early cell differentiation ^40^. While SMAD2,3 is described as being triggered mainly by TGF-βs and activins ^41^, it was also shown to be triggered by sBMPs ^42^. We decided to include it in our study and quantify the phosphorylation of the nuclear translocated SMAD2. One of the markers for the so-called non-canonical pathway is the induction of alkaline phosphatase (ALP) 40, which is often taken as later osteogenic readout ^33^.

We then studied the phosphorylation and nuclear translocation of SMAD1,5,9 and SMAD2 after 4-5 h of culture on the films with increasing BMP concentrations (**Fig. 2**, **Fig. SI 8**). In order to avoid differences in fluorescent signal intensity between experiments, we normalized the fluorescent signals of pSMAD1,5,9 and pSMAD2 of each experiment by the non-BMP condition (**Fig. 2A** and **D**). Not-normalized data is shown in **Fig. SI 8**. Representative examples of pSMAD fluorescent signal values distribution, together with representative pictures of C2C12 cells presenting low and high pSMAD signals, are shown in **Fig. SI 7**.

**FIGURE 2.**
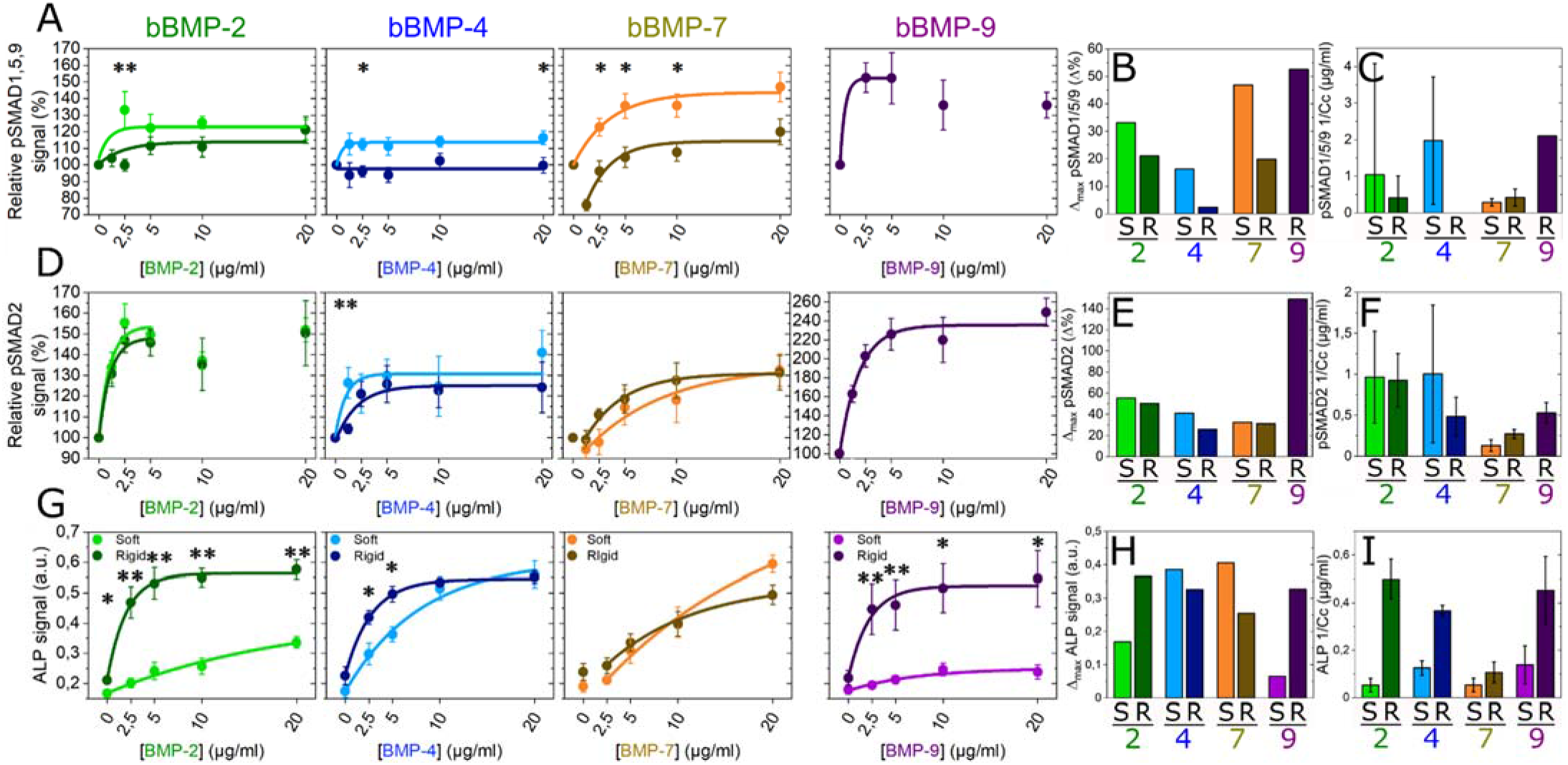
pSMAD and ALP analyses for the four bBMPs on soft and rigid films. Fluorescent signal quantification of (**a**) pSMAD1,5,9, (**d**) pSMAD2, and (**g**) ALP activity as a function of the BMP concentration in soft (light color) and rigid films (dark color). Quantitative parameters Δ_max_ and 1/Cc extracted from the fit of each of the experimental curves for (**b, c**) pSMAD1,5,9, (**e, f**) pSMAD2 and (**h, i**) ALP. Data represent the mean ± SEM of 2 to 4 independent experiments with two samples per condition in each independent experiment (at least 400 cells were analyzed in total). Statistical tests were done using non-parametric Kruskal-Wallis ANOVA test to compare S and R films (*p < 0.05; **p < 0.01).

After fitting the curves, as we did with cell adhesion, and extracting the parameters Δ_max_ and 1/Cc, we observed that, for pSMAD1,5,9 Δ_max_ value was higher on soft than on rigid films with bBMP-2, −4 and −7 (**Fig. 2B**). The sensitivity value (1/Cc) instead, was higher on soft than on rigid films with bBMP-2 and −4, but it was stiffness-independent for bBMP-7, which gave the lowest sensitivity. Nevertheless, attention should be paid due to big error bars (**Fig. 2C**). Curiously, at lower bBMP-7 concentrations on rigid films, the pSMAD1,5,9 values were below that of the non-BMP condition (**Fig. 2A** and **SI 8A**). Regarding bBMP-9, the cell response could solely be measured for cells cultured on stiff films, since C2C12 cells were barely adhering after 5 h on the soft films. bBMP-9 induced a high pSMAD1,5,9 response **(Fig. 2A** and **SI 8A**), with a high sensitivity (**Fig. 2C**). To note, pSMAD1,5,9 response to sBMPs was solely increased for cells on stiff films with sBMP-2 and 9 (**Fig. SI 9**).

pSMAD2 signal was stiffness-independent in all conditions except for cell sensitivity parameter (1/Cc) with bBMP-4, where soft films induced a higher sensitivity than rigid ones. However, the error bar in this condition was big (**Fig. 2F**). bBMP-9 triggered the strongest pSMAD2 signal, followed by bBMP-2 (**Fig. 2D-F** and **Fig. SI 8B**). bBMP-4 induced a higher pSMAD2 than pSMAD1,5,9 signal in terms of Δ_max_ on both film rigidities (**Fig. 2E**), and sensitivity only on rigid films (**Fig. 2F**). Similarly, to pSMAD1,5,9, at low concentrations bBMP-7 induced slightly smaller pSMAD2 signal values than the control, but on soft films instead (**Fig. 2D** and **Fig. SI 8B**).

The ALP response of C2C12 cells was quantified after 3 days of culture ^24^. The results corresponding to rigid films were extracted and adapted from P. Machillot *et al.* ^24^. Here, cells on soft films with bBMP-9 could be analyzed, since they could better adhere after 3 days of culture. bBMP-2 and bBMP-9 exhibited a strong stiffness-dependent response, with a significantly higher ALP activation on rigid films than on soft ones (**Fig. 2G-I** and **Fig. SI 10**). Conversely, the ALP response to bBMP-4 and −7 was very high for both soft and stiff films with only little stiffness-dependence (**Fig. 2**G-I). The ALP response to bBMP-4 and −7 on soft films was also higher than that to bBMP-2 and −9. Generally, for the ALP response, the sensitivity of cells to bBMPs was higher on rigid films (**Fig. 2I).**

Altogether, our data for C2C12 cells showed BMP-specific and stiffness-dependent responses. For bBMP-2, ALP response was stiffness-dependent, and to a less extend pSMAD1,5,9 signal. bBMP-4 induced a strong ALP response whatever the film stiffness, and to a less extend pSMAD1,5,9 and pSMAD2 signals. bBMP-7 also strongly activated ALP, to a lesser extent pSMAD1,5,9 and even less pSMAD2. Lastly, bBMP-9 induced the highest SMAD response from all the four BMPs, and a high ALP response.

### Involvement of BMP receptors and integrins in the cell adhesive response

Having evidenced that bBMPs can act as adhesive molecules to trigger myoblast and periosteum stem cell adhesion and spreading, we aimed to unravel the specific role of the BMP receptors and of integrins in the cell adhesive response. Firstly, an *in silico* screening was performed using RNA sequencing data from UCSC genome browser (Encode database). Thus, we identified the adhesion receptors expressed in C2C12 cells and expressed their relative abundance by quantifying the percentage (**Fig. SI 11**). We then verified the expression level of the BMP receptors (BMPR) and integrins of C2C12 myoblasts and hPDSCs using quantitative PCR (**Fig. 3A** and **B**). For C2C12 myoblasts, mRNA transcripts for the type I BMPR, ALK2, ALK3 and ALK5 were detected. ALK1, ALK4 and ALK6 were expressed at a much lower level. For the type II BMP receptors, BMPR-II, ACTR-IIA and to a much lower extent ACTR-IIB were detected. Regarding integrins, in both cell types β1 was the most highly expressed followed by β5 and β3. For hPDSCs, the similar BMP type I, II and integrins were detected, although their expression was quantitatively lower.

**FIGURE 3.**
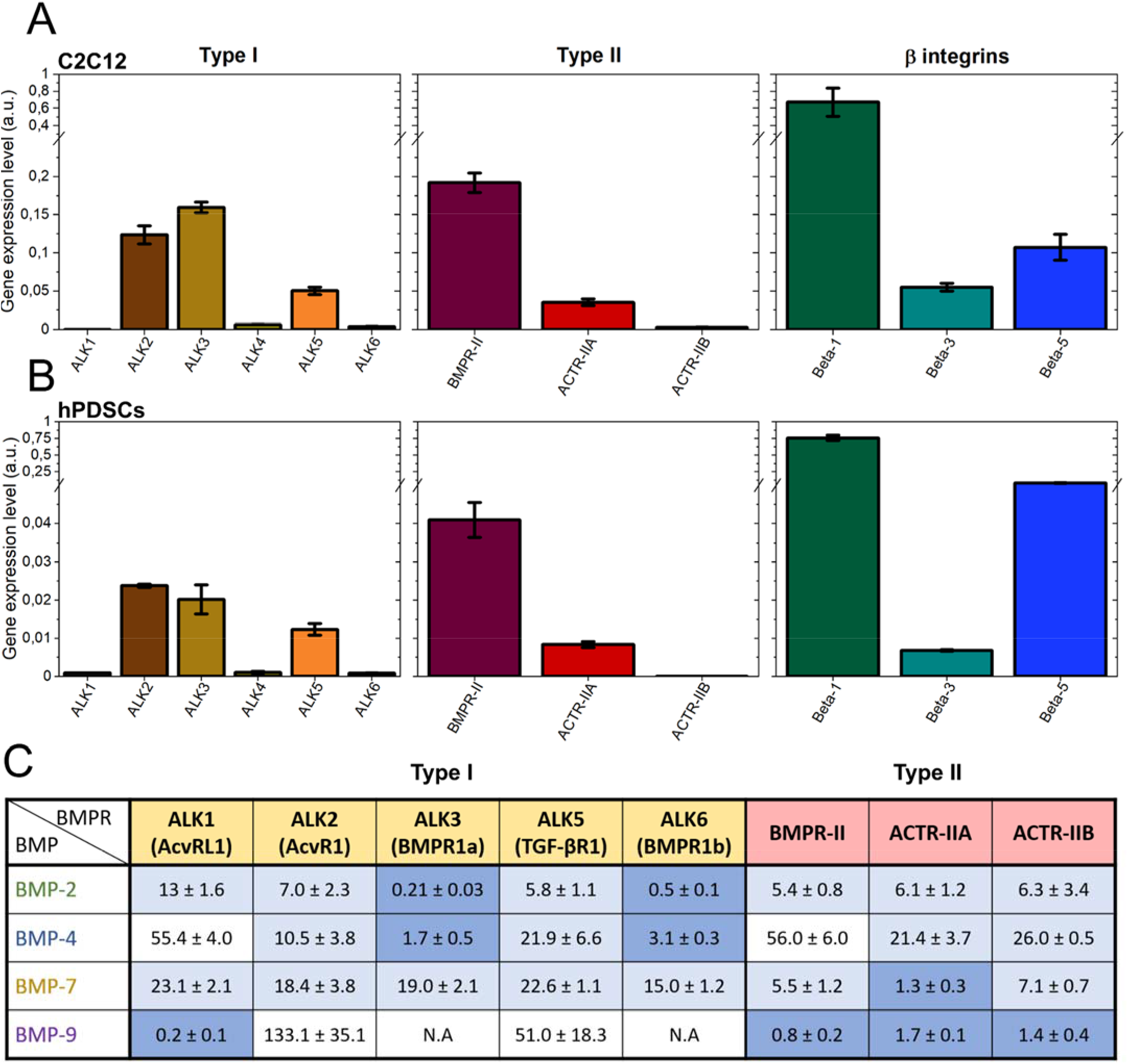
Relative gene expression analysis of type I, type II BMP receptors and β-integrins. Data corresponding to (**a**) C2C12 cells and to (**b**) hPDSCs. The gene expression was analyzed by qPCR and it was normalized to the expression of EF1, PPIA and GUSB genes. Data are represented as mean ± SEM of three independent experiments. (**c**) Table of the affinity constants (K_d_ in nM) of the type I and II BMP receptors with the four selected BMPs. Data was obtained from kinetic experiments performed by bio-layer interferometry (BLI), from the work of Khodr *et al.* ^43^. The highest affinity couples (Kd < 5 nM) are highlighted in dark blue, those with an affinity in the range from 5 to 25 nM are in light blue. N.A means no measurable interaction. Values are given as mean ± SD of three independent experiments.

Next, we quantified the molecular interactions between the four BMPs and the BMP receptors type I and II using reflectometric interference spectroscopy with the commercially-available biolayer interferometry setup. We obtained the affinity constants (Kd) values for each BMP-BMP receptor couple (**Fig. 3C**), as recently shown in Khodr *et al.* ^43^. Confirming the literature data ^4^, BMP-2 and BMP-4 exhibited high affinity for ALK3 and ALK6. BMP-7 exhibited a different behavior since it has the highest affinity for ACTR-IIA and has a moderate, non-specific affinity for all ALK receptors. BMP-9 had the strongest affinity for ALK1 and a similar affinity for the three type II receptors. We did not consider ALK4 for this study since it is an activin receptor and no evidence was found in the literature regarding a possible interaction with BMPs.

In order to assess the specific role of each receptor, type I (ALK2, 3, 5 and 6) and type II BMP receptors (BMPR-II, ACTR-IIA, ACTR-IIB), as well as β integrins (1, 3 and 5) were silenced using siRNA. The efficacy of the silencing was first verified (**Fig. SI 12**). The cell number and cell spreading area were analyzed on one particular BMP concentration (20μg/ml), normalized by the scrambled control condition and averaged, to reveal the contribution of each receptor (**Fig. 4** and **Fig. SI 13**). Non-normalized raw data are plotted in **Fig. SI 14**.

**FIGURE 4.**
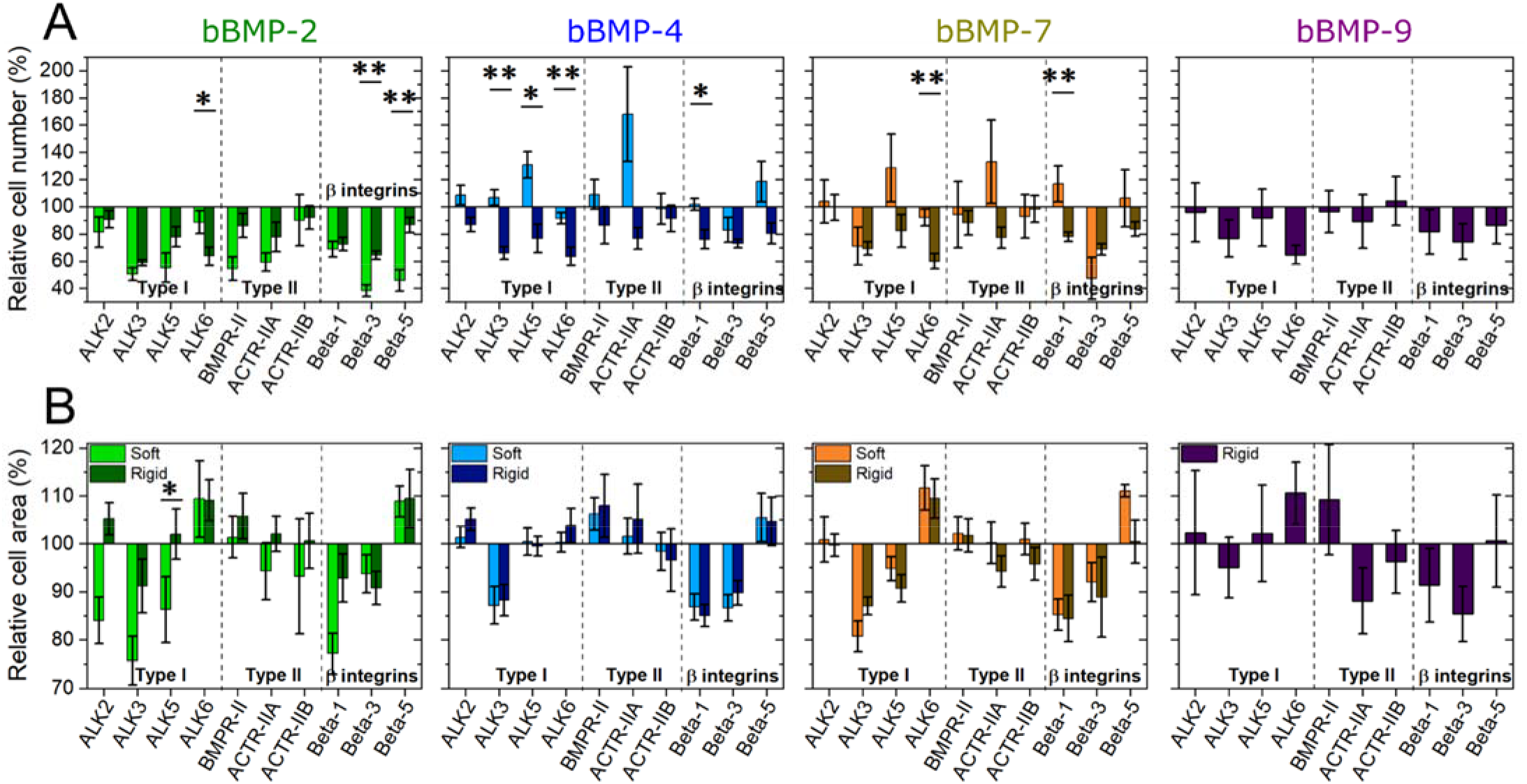
Effect of BMP receptor and β chain integrin silencing on cell adhesion and spreading. C2C12 cells were transfected with siRNA against BMP receptor type I (ALK2, ALK3, ALK5, ALK6), BMP receptor type II (BMPR-II, ACTR-IIA, ACTR-IIB) and β chain integrins (β1, β3, β5) and plated on soft (light color) and rigid (dark color) films with bBMPs for 5 or 4 h, respectively. The cell number par mm^2^ of substrate area and the spreading area were quantified. The relative % is given, in comparison to a control scrambled siRNA. (**a**) Relative cell number (%), (**b**) relative cell area (%). Data represent the mean ± SEM, with 3 biological replicates and 2 technical replicates per experiment. Statistical tests were done using non-parametric Kruskal-Wallis ANOVA test (*, p ≤ 0.05; **, p ≤ 0.01). Statistical comparisons were made between soft and rigid films for each condition.

Among the ALK receptors, on the non-BMP condition, ALK3 enhanced cell adhesion and spreading, while ALK6 promoted adhesion but inhibits spreading (**Fig. SI 13**). The same response was observed on films with bBMPs, with some particularities: ALK3 role on adhesion is stiffness-dependent with bBMP-4, and ALK6 induces adhesion only on rigid films but not on soft ones (**Fig. 4**). Notably, the response of ALK2 to bBMP-2 was mechano-sensitive regarding the cell spreading area. ALK5 was involved in a mechano-sensitive cell adhesion on bBMP-4 and −7, being an inhibitor on soft films and an activator on rigid ones (**Fig. 4A** and **Fig. SI 14A)**. It was also mechano-sensitive for cell spreading on bBMP-2, being activator on soft films and no role on rigid films (**Fig. 4B**).

Regarding type II BMP receptors, mainly BMPR-II and ACTR-IIA played a certain role in adhesion and spreading. On the non-BMP condition, both receptors induced adhesion, while BMPR-II inhibited spreading and ACTR-IIA activated it (**Fig. SI 13**). On films with bBMPs, BMPR-II was a slight inhibitor of cell spreading, but a strong activator of cell adhesion on soft films with bBMP-2. ACTR-IIA was mechano-sensitive, being an inhibitor of cell adhesion on soft films with bBMP-4 and −7, and activator on rigid films in response to all BMPs. It had a minor role on cell spreading, except for rigid films with bBMP-9 where it induced cell spreading (**Fig. 4**).

With respect to integrins, β3 integrin had an important and consistent role in activating both cell adhesion and spreading, independently of film stiffness and of the BMP presence. The role of β1 integrin was generally comparable to that of β3. β1 integrin was also involved in a mechano-sensitive response to bBMP-7, acting as an inhibitor of cell adhesion on soft films, and an activator on rigid ones. Lastly, β5 integrin was involved in a mechano-sensitive cell response to bBMP-2: it was an inhibitor of cell adhesion on soft films and no role on rigid ones. β5 integrin was an inhibitor of cell spreading on bBMP-2 and −4, on both film stiffness, and on soft films with bBMP-7 (**Fig. 4** and **Fig. SI 13**).

### Involvement of BMP receptors and integrins in early cell differentiation to bone

The effect of receptor silencing on early osteogenic differentiation was also investigated for pSMADs and ALP activity (**Fig. 5** and **Fig. SI 15**). Non-normalized raw data are plotted in **Fig. SI 16**.

**FIGURE 5.**
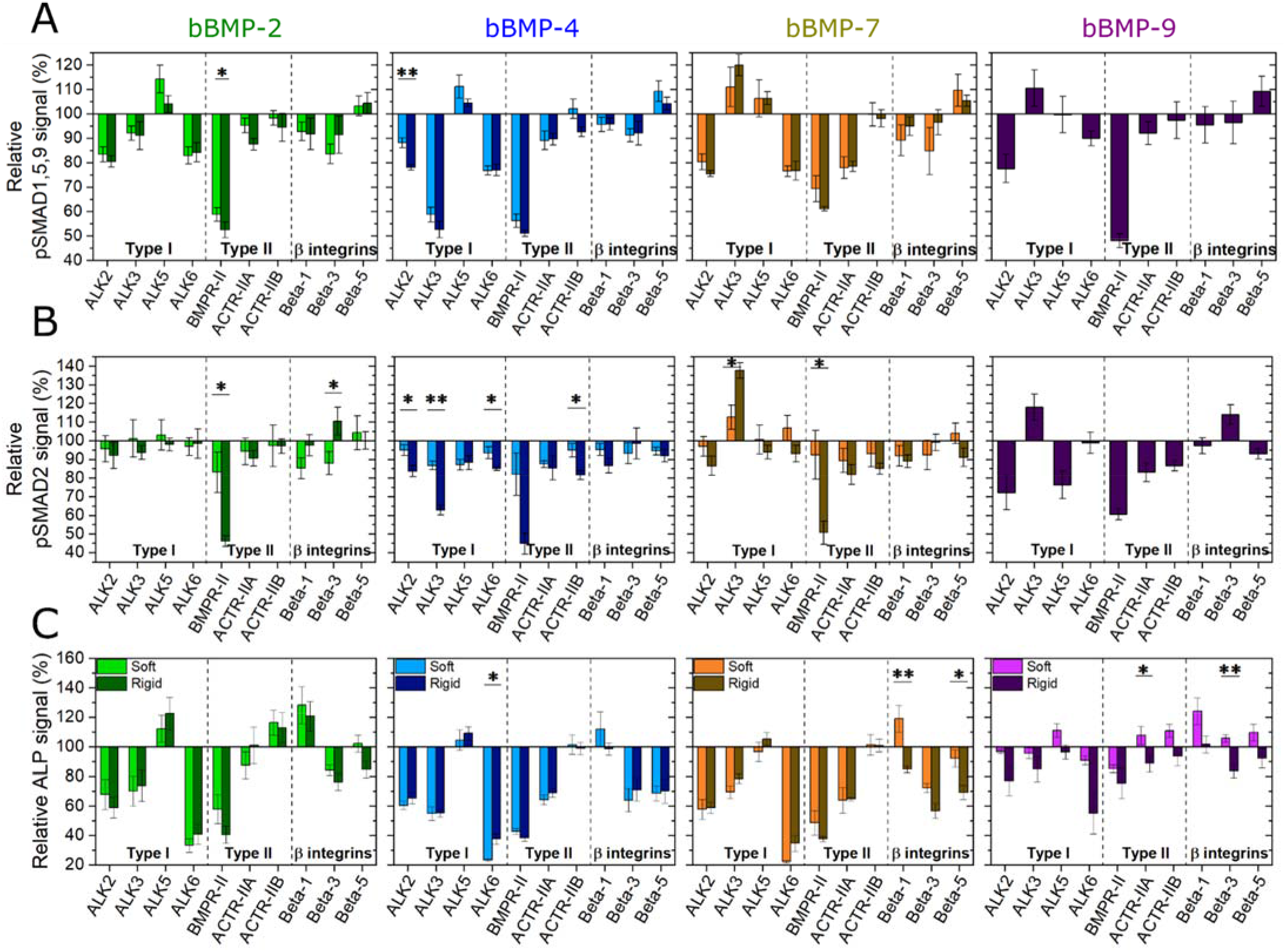
Effect of BMP receptor and β chain integrin silencing on early cell differentiation to bone. C2C12 cells were transfected with siRNA against BMP receptor type I (ALK2, ALK3, ALK5, ALK6), BMP receptor type II (BMPR-II, ACTR-IIA, ACTR-IIB) and β chain integrins (β1, β3, β5) and plated on soft (light color) and rigid (dark color) films with bBMPs for 5 or 4 h, respectively. (**a**) pSMAD1,5,9 signal and (**b**) ALP activity were quantified, and the relative % is given, in comparison to a control scrambled siRNA. Data represent the mean ± SEM. Statistical tests were done using non-parametric Kruskal-Wallis ANOVA test (*, p ≤ 0.05; **, p ≤ 0.01). Statistical comparisons were made between soft and rigid films for each condition.

Regarding pSMAD1,5,9, ALK2 exhibited an important role for all BMPs in a non-specific manner. ALK3 revealed to have a BMP-specific role: it is an activator for bBMP-4 and an inhibitor for bBMP-7 and −9. ALK6 was an activator of pSMAD1,5,9 on all bBMPs. ALK5 had only a minor inhibiting role for bBMP-2, −4 and −7, which was more visible on soft films than rigid ones (**Fig. 5A)**.

With respect to the BMPR type II receptors, BMPR-II was an important and a non-specific activator for all BMPs on both film stiffness, as well as for the no BMP condition (**Fig. 5A** and **Fig. SI 15A**), while ACTR-IIA was an activator of the pSMAD1,5,9 response to bBMP-7, independently of film stiffness.

Lastly, β integrins were found to have a minor role on SMAD1,5,9 activation in comparison to ALKs and BMP-type II receptors. β3 integrin was the most important activator, notably for bBMP-2 and −4, on both film stiffness, and solely on soft films for bBMP-7. β5 integrin had a minor inhibiting role on soft films with bBMP-4 and −7, and on rigid films with bBMP-9 (**Fig. 5A**).

For pSMAD2 (**Fig. 5B**), BMPR-II was again an important and non-specific activator for all bBMPs, and the no BMP condition (**Fig. SI 15**), especially on rigid films. Interestingly, here again, ALK3 had a notable role: being a stiffness-dependent activator for the pSMAD2 response to bBMP-4, a stiffness-dependent inhibitor for the response to bBMP-7, and an inhibitor for bBMP-9, its role being prominent on rigid films in comparison to soft ones.

Regarding the ALP activity (**Fig. 5C** and **Fig. SI 16, 17**), generally, ALK2, ALK3 and ALK6 were activators, whereas ALK5 had only an inhibitor role for bBMP-2 (**Fig. 5C**). BMPR-II and ALK6 played the most important role in activating ALP. However, ALK6 was a stronger ALP activator than BMPR-II on bBMP-2, −4 and −7 for soft films, and on bBMP-9 for rigid ones. ACTR-IIA played a role in ALP activation on bBMP-4 and −7, independently of film stiffness. ACTR-IIB only played a minor role inhibiting ALP on bBMP-2 (**Fig. 5C**). Regarding integrins, β3 and β5 integrins were generally activators of ALP for bBMP-2, −4 and −7. β1 integrin was rather inhibiting ALP notably on soft films and had almost no effect on stiff films, except for BMP-2 (**Fig. 5C**).

Globally, the role of the receptors was related to each readout of the osteogenic differentiation (pSMAD1,5,9; pSMAD2 or ALP) with only little differences depending on the film stiffness, and little differences between the bBMPs except for the ALK3 and ACTR-IIA receptors.

## Discussion

### Stiffness dependence of BMPs and BMP receptors in cell adhesion and spreading

For the first time, the effect of 4 different BMPs, presented in a matrix-bound manner, has been compared on cell adhesion, spreading, SMAD signaling and ALP activity. We show that generally, film stiffness plays a relevant role on the BMP-induced cell responses. Furthermore, we show the role of BMPR type I, type II, and three β-chain integrins, on the aforementioned cell responses for each condition. This study was feasible thanks to a high-content automatized procedure for film preparation ^24^, image acquisition and analysis

We found that all four studied bBMPs are able to induce both cell adhesion and cell spreading (**Fig. 2**, **Fig. 6** and **Fig. SI 2, 4** and **5**). This extends our initial findings regarding the role of bBMP-2 on cell adhesion and migration ^23^. This effect of BMPs on cell adhesion and spreading was revealed thanks to the matrix-bound presentation of the BMPs together with the variation of film stiffness. Among the four BMPs, bBMP-4 and bBMP-7 induced the most potent adhesion and spreading response, on C2C12 and hPDSCs (**Fig. 1D-G** and **Fig. SI 4D-G**). All four bBMPs have a stronger role inducing cell spreading on rigid films than on soft ones (**Fig. 2F** and **G**). However, on the no BMP condition, only cell number is increased on rigid films with respect to soft ones, but not cell spreading, as it was found in previous works ^23,26^. bBMP-9 on soft films did not induce neither cell adhesion nor cell spreading on C2C12 (**Fig. 1B** and **C**), and only a small increase is observed on hPDSCs (**Fig. SI 4B** and **D**). The fact that BMP-9 does not bind to ALK3 and ALK6 could help explaining this observation, since these two receptors are important for cell adhesion (**Fig. 4A**), and ALK3 also for cell spreading (**Fig. 4B**). Our data are consistent with other works showing a positive effect of BMP-2, −4 and −7 on cell adhesion and migration with other cell types ^26,30,44^

**FIGURE 6.**
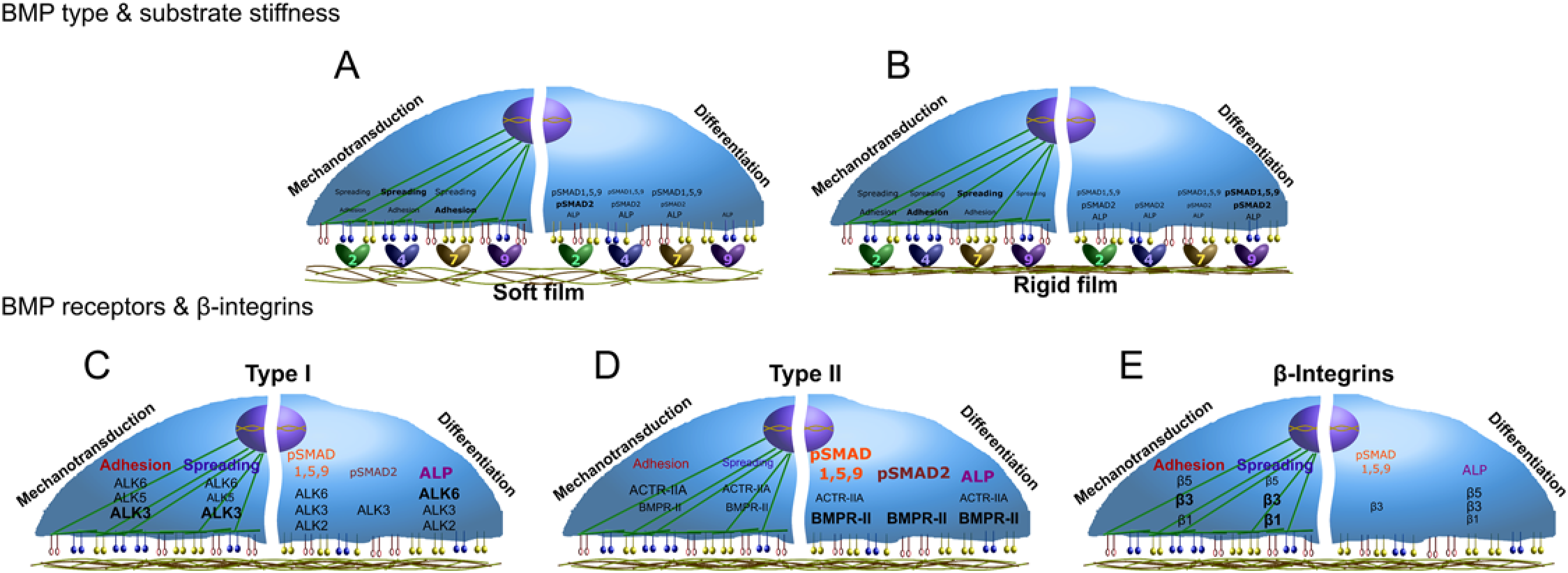
Schematic representations summarizing the major results of this study. The influence of the 4 bBMPs, depending on the film stiffness, on C2C12 cell adhesion and spreading, as mechanotransduction parameters studied, and pSMAD1,5,9, pSMAD2 and ALP activity, as differentiation parameters studied, is represented for (**a**) soft and (**b**) rigid films. Moreover, the role of the BMP receptors type I, type II and the three β chain integrins analyzed, on the mechanotransduction and differentiation parameters aforementioned, are depicted for each type of receptor: (**c**) type I BMPR, (**d**) type II BMPR, and (**e**) β chain integrins. Since no relevant stiffness-dependent differences were generally observed for the receptors, no specific detailed distinction is represented.

ALK2 appears to be mechano-sensitive in response to bBMP-2 only (**Fig. 4**), by inducing cell spreading on soft films. Our data are consistent with the recently reported mechanosensitivity of ALK2 ^45^. Interestingly, this mechanosensing appears to be altered during heterotypic ossification associated to fibrodysplasia ossificans progressiva (FOP).

ALK3 appears to be a central receptor in cell adhesion and spreading for all BMPs (**Fig. 4**).. ALK3 has a major role in several cancers ^5^ and plays a role in the adhesion of human epithelial ovarian cancer spheroids to the substratum ^46^.

We found that ALK5 tends to be mechano-sensitive for cell spreading on bBMP-2, and for cell adhesion on bBMP-4 and 7 (**Fig. 4**). Our data are consistent with the role of ALK5 described in other contexts showing a mechanosensing role of this receptor. In fact, ALK5 is historically known as a TGF-β receptor ^47^ and for its involvement in EMT in breast epithelial ^48^ and endothelial cells ^49^. It was already shown to be involved in the response to shear stress in the endothelium ^50^, in chondrocyte mechanosensing ^51^ and in the activation of focal adhesion kinase (FAK) in lung epithelial cells ^52^.

Our data revealed an important mechano-sensitive role of ACTR-IIA in cell adhesion, notably in response to both bBMP-4 and bBMP-7 (**Fig. 4A**). This finding is in line with a work showing a mechano-sensitive role of ACTR-IIA in tenogenic commitment of adipose stem cells under magnetic stimulation ^53^.

### Identification of integrins in BMP-induced cell adhesion and BMP signaling

Regarding the role of integrins, β3 integrin appears to have a major role in cell adhesion and spreading on all bBMPs. The role of β3 integrins is crucial for cell adhesion in soft environment, which corresponds to our finding, whereas both β1 and β3 integrins cooperate in a stiff environment as already described ^54^. β1 integrin was also important for cell adhesion on all bBMPs, with a peculiar stiffness-dependent response for cell adhesion to bBMP-7. β5 integrin also exhibited a stiffness-dependent response to bBMP-2 and −4 (**Fig. 4**).

Apart from their role in adhesion, integrins are also important for bone differentiation ^55^. In our experimental conditions, we found that β1 integrin is a slight inhibitor of BMP-2-mediated C2C12 ALP activity for soft and stiff films, while it was stiffness-dependent on bBMP-7 (**Fig. 5C**). The role of β1 integrin in osteoblastic differentiation is recognized, but appears to depend on the cell type and on the context. Previously in our group, Sefkow-Werner *et al.* found an activator role of β1 integrin with sBMP-2 and BMP-2 bound to heparan sulphate, by using also C2C12 cells ^27^. Many data have shown that β1 integrin plays an important role in osteoblast differentiation and function ^56–58^. Mice expressing a dominant-negative β1 integrin in mature osteoblasts show reduced bone mass and defective bone formation ^56^. β5 integrin has been shown to be involved in bone differentiation in a non-Smad-dependent manner ^59^.

### Stiffness dependence of BMP and BMP receptors in SMAD and ALP signaling

BMPs are traditionally referred to activate the pSMAD1,5,9 pathway ^41^ whereas TGF-β activates SMAD2,3 signaling ^47^. To our knowledge, to date, there was no systematic comparison of the role of BMPs on SMAD and ALP signaling available in the literature. Here, we showed that all four bBMPs are able to activate both pSMAD1,5,9, pSMAD2 and ALP pathways in a BMP-specific and, in certain conditions, in a stiffness-dependent manner (**Fig. 6A** and **B**). In the stiffness-dependent conditions, pSMAD1,5,9 and pSMAD2 Δ_max_ values were always higher on soft films (**Fig. 2A, B, D**, **and** **E**). Notably, bBMP-4 induced a negligible pSMAD1,5,9 response on stiff films while bBMP-9 induced the highest one.

BMP-2 was recently shown to promote both canonical SMAD1,5,8 and non-canonical SMAD2,3 pathways in trophoblast cells ^60^. Soluble BMP-4 also activates SMAD3 ^61^. Our data is also consistent with SMAD and non-SMAD signaling reported for BMP-7 induced BMP signaling in tenocyte-like cells ^62^.

Regarding pSMAD1,5,9 signaling, ALK2 and ALK6 were found to have a central role independently of the BMP type, while the effect of ALK3 was bBMP-specific: it is a strong activator on bBMP-4 and inhibitor on bBMP-7 and −9 (**Fig. 5A**).

BMPR-II had a major activator role for all bBMPs while ACTR-IIA was specifically associated to the response to bBMP-7 (**Fig. 5A**). Our data are consistent with the role of ACTR-IIA for BMP7-mediated chemotaxis of monocytes and pSMAD activation ^63^

The important role of bBMP-9 activating SMAD1,5,9 and SMAD2, could be correlated with the strong binding to BMPR-II (**Fig. 3**), which plays a crucial role in SMAD activation (**Fig. 5A** and **D**), and to ALK1 (**Fig. 3**) which is also important for SMAD2,3 and SMAD1,5 activation ^64^. Also, the fact that BMP-9 is resistant to the BMP inhibitor Noggin may help to the strong SMAD activation ^65^.

ALP is a common marker to assess BMP-induced osteoblastic differentiation ^33^. We showed that ALP is highly expressed in C2C12 cells in response to bBMPs (**Fig. 2G** and **Fig. SI 10**). Our data are in agreement with the previously demonstrated ALP activities evidenced for soluble BMP-2, −4, −7, and −9 ^33^. We showed here for the first time a BMP-specific stiffness-dependent ALP response of cells cultured on bBMPs. Indeed, ALP was stiffness-dependent in response to bBMP-2 and −9 and stiffness-independent for bBMP-4 and −7 (**Fig. 2G-I**). The relevant roles of ALK2 and ALK3 in ALP activation that we observed, are consistent with role of ALK2 in ALP activity ^66^ as well as the role of ALK3 ^67^, found in the literature. Our data also showed that ACTR-IIA is an activator of BMP-4 and 7-mediated ALP activity, while ACTR-IIB was an inhibitor solely for the ALP response to bBMP-2 (**Fig. 5C**). They are consistent with the role of ACTR-IIA and B receptors in the regulation of bone mass ^68^.

In conclusion, in this work we carried out an extensive study where we shed light on the effect of 4 matrix-bound BMPs combined with 2 film rigidities (soft and rigid), on early phases of cell mechanotransduction, with C2C12 cells and hPDSCs, and osteogenic differentiation, with C2C12 cells. Moreover, by means of siRNA, we revealed the role of type I and type II BMP receptors as well as 3 beta chain integrins, on the aforementioned cell processes. This study paves the way for the elucidation of the complex interplay between BMPs, substrate physico-chemical properties, BMP receptors and integrins, on the cell fate towards osteogenic lineage, which is crucial for the development of new, tailored, medical implants.

## Methods

### Polyelectrolyte film buildup, crosslinking and BMPs loading

Poly(L-lysine) hydrobromide (PLL) and poly(ethylene imine) (PEI) were purchased at Sigma-Alrich (St Quentin Fallavier, France), and hyaluronic acid (HA) at Lifecore medical (USA). PLL and HA were dissolved in a HEPES-NaCl buffer (20 mM Hepes and 0.15 M NaCl at pH 7.4) at 0.5 mg/mL. PEI was dissolved in a NaCl 0.15 M solution at pH ~6.5, at 4 mg/mL. A first layer of PEI was always deposited. All rinsing steps were performed with 0.15 M NaCl at pH ~6.5. LbL films were directly deposited in 96-well cell culture microplates (Greiner bio-one, Germany) using an automated liquid handling robot (TECAN Freedom EVO® 100, Tecan France, Lyon) and a custom-made macro ^24^. The films were then chemically crosslinked using 1-Ethyl-3-(3-Dimethylamino-propyl)Carbodiimide (EDC) at a final concentration of 30 or 70 mg/mL, and N-Hydrosulfosuccinimide sodium salt (Sulfo-NHS) at a concentration of 11 mg/mL as catalyzer, as previously described ^38,69^. BMP loading in the films was done following an established protocol ^70^. BMP-2 was purchased from Bioventus (France), BMP-4 and −9 from Peprotech (France) and BMP-7 from Olympus Biotech (France).

### Cell culture, and quantification of stem cell adhesion and differentiation

We used two types of BMP-responsive cells, skeletal myoblasts ^70^ and periosteum-derived stem cells (hPDSCs) ^37^ to assess the bioactivity of the biomimetic coatings at high content. C2C12 skeletal myoblasts (<15 passages, obtained from the American Type Culture Collection, ATCC) and hPDSC (<12 passages, obtained from F. Luyten) were cultured as previously described ^37,70^ in tissue culture flasks, in a 1:1 Dulbecco’s Modified Eagle Medium (DMEM):Ham’s F12 medium (Gibco, Life Technologies, France) for C2C12 and in high-glucose Dulbecco’s modified Eagle’s medium (high DMEM) + pyruvate for hPDSC (Gibco, Life Technologies, France), supplemented with 10% fetal bovine serum (FBS, Gibco, Life TechnologiesFrance) and 1% penicillin/streptomycin (Gibco, Life Technologies, France) in a 37 °C, 5% CO_2_ incubator. For analysis of cell adhesion and pSMAD, 8300 cells/cm^2^ of C2C12 and 7000 cells/cm^2^ of hPDSCs were seeded in each well. For analysis of ALP, 33000 cells/cm^2^ of C2C12 were seeded in each well. For quantification of cell adhesion, cells were cultured for 4 or 5 h and then fixed with 4% paraformaldehyde. Nuclei were stained with 4′,6-diamidino-2-phenylindole (DAPI) (Invitrogen, Thermofisher Scientific, France) and the actin cytoskeleton with Alexa Fluor 633-phalloidin (Thermofisher Scientific, France). For quantification of cell differentiation, C2C12 cells were stained at 4 or 5 h for pSMAD analysis, and after 3 days of culture for ALP analysis. pSMAD was stained by using a 1:400 dilution of an antibody anti-pSMAD1,5,9 (Cell Signaling, ref: 13820S) or a 1:800 dilution of an antibody anti-pSMAD2 (Cell Signaling, ref: 18338S) both diluted in a solution of bovine serum albumin (BSA, Sigma-Aldrich, Merck, Germany) 3% w/v in PBS. A secondary antibody conjugated to Alexa Fluor 555 (Invitrogen, Thermofisher Scientific, France) diluted at 1:500 in BSA 3% w/v in PBS, was used for both SMAD markers since they were always analyzed in different samples. ALP was stained by using fast blue RR salt (Sigma-Aldrich, Merck, Germany) in a 0.01 % (w/v) naphtol AS-MX solution (Sigma Aldrich, Merck, Germany), according to the manufacturer’s instructions. ALP was imaged using a scanner and quantified using a TECAN Infinite 1000 microplate reader (TECAN France, Lyon) by measuring the absorbance at 570 nm using the multiple-read/well mode ^24^. The mean value of 76 measured different positions per microwell was taken.

For the quantification of immunofluorescence stainings, a GE INCA 2500 imaging system (General Electrics Healthcare, France) was used with an inverted 20X objective (Plan Apo N.A.=0.75, Nikon, Japan). 21 images distributed over the well were acquired and, generally, at least 100 cells were analyzed par well and plate. Between 2 and 4 independent experiments (biological replicates) were performed with 2 technical replicates per experiment, making up a total of, generally, at least 400 cells per condition. For comparison of pSMAD1,5,9 and pSMAD2 fluorescent signal, the same exposure time was always used for each marker.

For image analysis, InCarta software (General Electrics Healthcare, USA) was used to automatically segment nuclei and cells, and to quantify cell number, cell area and pSMAD signal intensity.

For quantitative analysis of the BMP dose-response experiments, the data were fitted with an exponential function:

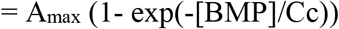

Where *Cc* is the characteristic concentration of cell adhesion, and A_max_ is the plateau value (in number of cells/mm^2^ for the cell number, and in μm^2^ for the cell area). In some cases where the curve decayed after reaching a maximum, only the ascending part of the curve was fitted.

In the BMP dose-response experiments, Δ_max_ was calculated by subtracting the maximum value by the value corresponding to the control condition without BMP.

### Expression of BMP and integrin receptors

Expression of BMP type I, type II and integrin receptors was investigated by mRNA transcript analysis. Total RNA from expanded C2C12 and hPDSCs was extracted by lysing cells in lysis buffer supplemented with β-mercaptoethanol. Further RNA extraction was performed using the RNeasy kit (Qiagen) according to the manufacturer’s instructions. Complementary DNA (cDNA) was obtained by reverse transcription of total RNA with Oligo (dT)20 as primer (Superscript III; Invitrogen, Thermofisher Scientific, France). Quantitative PCR was performed in duplicates with a thermocycler (MX4800P; Agilent Technologies, France) as previously described ^26^ using the SYBER green kit. The primer sequences are given in **Table SI 2**. Primer efficiency was established by a standard curve using sequential dilutions of gene-specific PCR fragments. Data were normalized from the quantitative RT-PCR housekeeping genes EF1, Gusb, and PPIA as an index of cDNA content after reverse transcription.

### SiRNA interference

Cells were transfected with siRNA against β1, β3 or β5 integrins, BMP receptors type I (ALK2, ALK3, ALK5, ALK6), BMP receptors type II (BMPR-II, ACTR-IIA, ACTR-IIB) (ON-TARGET plus SMART pool; respectively, mouse ITGB1, ITGB3, ITGB5, ALK2, ALK3, ALK5, ALK6, BMPR-II, ACTR-IIA, ACTR-IIB) (Dharmacon, France). The gene target siRNA sequences used for transfection are listed in **Table SI 3**. A scrambled siRNA (Dharmacon, France) was taken as control. Cells were seeded at 5,000 cells/cm^2^ in 12-well plates and cultured in 1 ml of GM with 0.5% of antibiotics for 24 h. The transfection mix was prepared as follows: For one well, 2.8 μl of Lipofectamine RNAiMAX Reagent (Invitrogen, Thermofisher Scientific, France) was added to 60 μl of Opti-MEM medium (Gibco, Life Technologies, France) and 1.2 μl of 16.67 μM siRNA, diluted in RNAse free water (5 Prime) was added to another 60 μl Opti-MEM medium. Lipofectamine-containing mix was added to siRNA-containing mix and incubated for 20 min at RT. Then, 120 μl of the final mix was added to each well. After 24 h of incubation at 37°C, the cells were transfected for the second time and incubated for another 24 h. The cells were then detached with 0.25% trypsin-EDTA (Gibco, Life Technologies, France), seeded in GM on the films, and allowed to adhere.

### Data representation and statistical analysis

For box plots, the box shows 25, 50 and 75% percentiles, the square shows the mean value and the error bars correspond to the standard deviation. For scatter plots and bar plots, the mean values and the standard error of the mean (SEM) are represented. Experiments were performed in duplicate, triplicate, or quadruplicate (biological replicates) with 2 wells per condition (technical replicate) in each experiment. Statistical analysis was performed using the non-parametric analysis Kruskal-Wallis analysis of variance (K-W ANOVA) to obtain *p* values (*p* < 0.05 was considered significant).

In the adhesion experiments, the mean values of cell number and cell area were obtained by averaging all the values for all the wells (technical and biological replicates). To calculate the mean value of pSMAD signal, first the average of all values per well was calculated, then the average of all wells from all the experiments was calculated. In the ALP experiments, the mean value was obtained by averaging the mean absorbance values of each well from all the experiments. In the knockdown experiments, the relative values were calculated by first averaging all the mean values per well from all the experiments, then normalizing the values by the scrambled condition.

## Acknowledgements

C. Picart is a senior member of the Institut Universitaire de France, whose financial support is acknowledged. The study was supported by Agence Nationale de la Recherche (ANR CODECIDE, ANR-17-CE13-022 to C.A.R. and C.P, ANR GlyCON, ANR-19-CE13-0031-01 E.M.), by the Fondation Recherche Medicale (FRM) (grant DEQ20170336746 to C.P. DEQ20170336702 to C.A.R), by the European Research Council (ERC Biomim, GA 259370, POC BioactiveCoatings, GA692924). We are part of the GDR2088 “BIOMIM” and the GDR “Réparer l’humain” research networks. V.K. was supported by a PhD fellowship from Grenoble Institute of Technology. A. Guevara-Garcia is supported by a CONACYT fellowship from the mexican government (CVU: 532484). We thanks Prof. Franz Luyten for providing the hPDSCs cells and for fruitful discussions. We are grateful to David Pointu for his technical advices.

## Author contribution

C.P. supervised the project, designed the experiments, wrote the manuscript, acquired funding and provided the resources. A.S and P.M. designed, and performed experiments, analyzed the data, and wrote the manuscript. V.K. carried out experiments, analyzed the data and edited the manuscript. L.F. and C.A.R provided advice and technical assistance, and edited the manuscript. E.M. and A.G.G reviewed and edited the manuscript.

## Materials & Correspondence

Catherine Picart and Adrià Sales

