## Supplemental File for "Differential bioactivity of four BMP-family members as function of biomaterial stiffness"

**TABLE SI 1.** Percentage of BMP proteins (BMP-2, -4, -7 and -9) incorporated in the $(PLL/HA)_{12}$ films and the corresponding estimated density on the film. The samples analyzed correspond to an initial BMP loading concentration in solution of 20 µg/mL. The amount of BMP was quantified using a micro bicinchoninic acid assay (microBCA) assay. Experiments were performed at least in triplicate (data are mean ± SD of 2to 3 independent experiments).

| BMP type | % of BMPs incorporated | | BMPs surface densities (µg/cm²) | |
|---|---|---|---|---|
| | Soft film | Rigid film | Soft film | Rigid film |
| **BMP-2** | 60 ± 3 % | 65 ± 5 % | 1.81 ± 0.09 | 1.97 ± 0.15 |
| **BMP-4** | 82 ± 4 % | 67 ± 9 % | 2.48 ± 0.12 | 2.03 ± 0.27 |
| **BMP-7** | 94 ± 3 % | 85 ± 8 % | 2.85 ± 0.09 | 2.58 ± 0.24 |
| **BMP-9** | 92 ± 1 % | 88 ± 1 % | 2.79 ± 0.03 | 2.67 ± 0.03 |

**TABLE SI 2**. Primer sequences.

| Mouse Gene | Forward sequence (5'-3') | Reverse sequence (5'-3') |
|---|---|---|
| ALK1 | GACACCCACCATCCCTAACC | TTGGGGTACCAGCACTCTCT |
| ALK2 | AGTGGAAGTCTGGAAAAGGAACA | TTCTCCCAGCAGGCTCCCTAT |
| ALK3 | GGAAGCATAGGTCAAAGCTGTTC | TGAGGGAGGCTTCCTTACAGA |
| ALK4 | ACGAGACAATCAACATGAAGCA | CGGAGGGCACTAAGTCGTAA |
| ALK5 | TTTCAGAGGGCACCACCTTA | AATGGTCCTGGCAATTGTTCT |
| ALK6 | CAACCCGGCCATAAGTGAAG | CTCCTTCTTGGTGCCCACATT |
| BMPR-II | CCCTCCCTTGACCTGGATAAC | TACAGCAACTGGACGCTCAT |
| ACTR-IIA | GCGTTCGCCGTCTTTCTTAT | AGCAAGGTTCAACACCAGTCT |
| ACTR-IIB | AGGGAAGCCTCCTGGGGATA | CCATGGCGTACATGTCGATAC |
| β1 | CGGACGCTGCGAAAAGATGA | CACATCGTGCAGAAGTAGGC |
| β3 | CCACACGAGGCGTGAACTC | CTTCAGGTTACATCGGGGTGA |
| β5 | CCCGTTATGAAATGGCCTCA | GCCTAGCTAGCGTGAGCAAA |
| EF1 | CCGTCAGAACGCAGGTGTTG | GTTCGCTTGTCGATTCCACC |
| PPIA | GTCTCCTTCGAGCTGTTTGC | GCGTGTAAAGTCACCACCCT |
| GUSB | CGGGACTTTATTGGCTGGGT | CCATTCACCCACACAACTGC |

| Human Gene | Forward sequence (5'-3') | Reverse sequence (5'-3') |
|---|---|---|
| ALK1 | TGGTGCTGTGGGAGATTGC | TCCTCAAAGCTGGGGTCATT |
| ALK2 | ATGTGACCAAGAGCCTGCAT | CGCAGGAGAGACCTTCACAC |
| ALK3 | GGTAGTGGGTCTGGACTACCT | ACGCCATTTGCCCATCCATA |
| ALK4 | CTCCTCCTTCTTCCCCCTTGTT | CCATCTGTCTCACACGTGTAGTTG |
| ALK5 | TTGCTGCAATCAGGACCATTG | AGATGCAGACGAAGCACACT |
| ALK6 | ATGACTCTGGGTTGCCTGTG | TCAATGGAGGCAGTGTAGGG |
| BMPR-II | TGAGCCCAACAGTCAATCCA | TGGCACACGCCTATTATGTGA |
| ACTR-IIA | GGCGTTTGCCGTCTTTCTTA | AACACGGTTCAACACCAGTTT |
| ACTR-IIB | GGGCCACAAGCCGTCTATT | GGAGGTTTCCCTGGCTCAAA |
| β1 | CCGCGCGGAAAAGATGAAT | CACAATTTGGCCCTGCTTGTA |
| β3 | TGCGAGTGTGACGACTTCTC | GTCAGTACGCGTGGTACAGT |
| β5 | TGCTTCGAGAGCGAGTTTGG | GTCCCCGATGTAACCTGCAT |
| EF1 | ATCCACCTTTGGGTCGCTTT | CTGAGCTTTCTGGGCAGACT |
| PPIA | TCAACCCCACCGTGTTCTTC | TGCTGTCTTTGGGACCTTGT |
| GUSB | GTGCGTAGGGACAAGAACCA | GGGAGGGGTCCAAGGATTTG |

**TABLE SI 3.** Gene target siRNA sequences used for transfection.

| Gene Target | Reference DHARMACON | siRNA target sequence (5' to 3') |
|---|---|---|
| β1 integrin | L-040783-01-0005, ON-TARGETplus Mouse Itgb1 (16412) - SMARTpool | UGCCAAAUCUUGCGGAGAA<br>UUACAAGAGUGCCGUGACA<br>GUGAAGACAUGGACGCUUA<br>CAAUGAAGCUAUCGUGCAU |
| β3 integrin | L-040746-01-0005, ON-TARGETplus Mouse Itgb3 (16416) – SMARTpool | AAACAGAGCGUGUCCCGUA<br>AAACACGUGCUGACGCUAA<br>GAGCAGUCUUUCACUAUCA<br>GUGAAAGAGCUGACGGAUA |
| β5 integrin | L-042453-01-0005, ON-TARGETplus Mouse Itgb5 (16419) - SMARTpool | CCGCUUAGGUUUCGGGUCU<br>GCUAGGCACGCACGGAUAA<br>AGAAGAUCGGAUGGCGAAA<br>ACUGCUAAGGACUGCGUUA |
| ALK2 | L-042047-00-0005, ON-TARGETplus Mouse Acvr1 (11477) - SMARTpool | CUAGAUCACUCGUGUACAU<br>GAAAUGGGAUCGUUGUAUG<br>UAUAAGAGGGUCGAUAUUU<br>GAAGGGCUGCUUUCAGGUU |
| ALK3 | L-040598-00-0005, ON-TARGETplus Mouse Bmpria (12166) - SMARTpool | GAGGAAUCGUGGAGGAAUA<br>GCUAGCUGGUUUAGAGAAA<br>GAAAUGGCUCGUCGUUGUA<br>GGCCAUUGCUUUGCCAUUA |
| ALK5 | L-040617-00-0005, ON-TARGETplus Mouse Tgfbr1 (21812) - SMARTpool | GGGCAGUUACUACAACAUA<br>CUAGAUCGCCCUUUCAUUU<br>GCGAAGGCAUUACAGUGUU<br>UGACAGCUUUGCGAAUUAA |
| ALK6 | L-051071-00-0005, ON-TARGETplus Mouse Bmpr1b (12167) - SMARTpool | GACAAUAGCUAAGCAAAUU<br>GGAAUGAGUGUAAUAAAGA<br>GCACAGAUGGGUACUGCUU<br>GACGAGAGCUUGAAUAGAA |
| BMPR-II | L-040599-00-0005, ON-TARGETplus Mouse Bmpr2 (12168) - SMARTpool | GCACAUAGGUCCCAAGAAA<br>GAACGCAACCUGUCACAUA<br>GCAUGAACCUUUACUGAGA<br>CUAAVAAGCUAGAUCCAAA |
| ACTR-IIA | L-040676-00-0005, ON-TARGETplus Mouse Acvr2a (11480) - SMARTpool | GAACCUUGCUAUGGUGAUA<br>GGACGCAUUUCUGAGGAUA<br>CAGACUUUCUUAAGGCUAA<br>GCAAUGCUCUGUGAAACGA |
| ACTR-IIB | L-040629-00-0005, ON-TARGETplus Mouse Acvr2b (11481) - SMARTpool | GCCCAGAAGUCACGUACGA<br>CGGCCUGGCUGUUCGGUUU<br>GAAGAGCGGGUAUCCCUGA<br>GGAACGAACUGUGCCACGU |
| Scrambled | D-001810-01-05) ON-TARGETplus Non-targeting | UGGUUUACAUGUCGACUAA |

**FIGURE SI 1. Representative fluorescent pictures of C2C12 cell adhesion and spreading on the films with matrix-bound BMPs.** C2C12 cells were fixed after 5h of culture on soft films, and after 4 h of culture on rigid films. Cells were stained for actin (in far-red) and nuclei (in blue) for increasing loading concentrations. Scale bar=100 µm

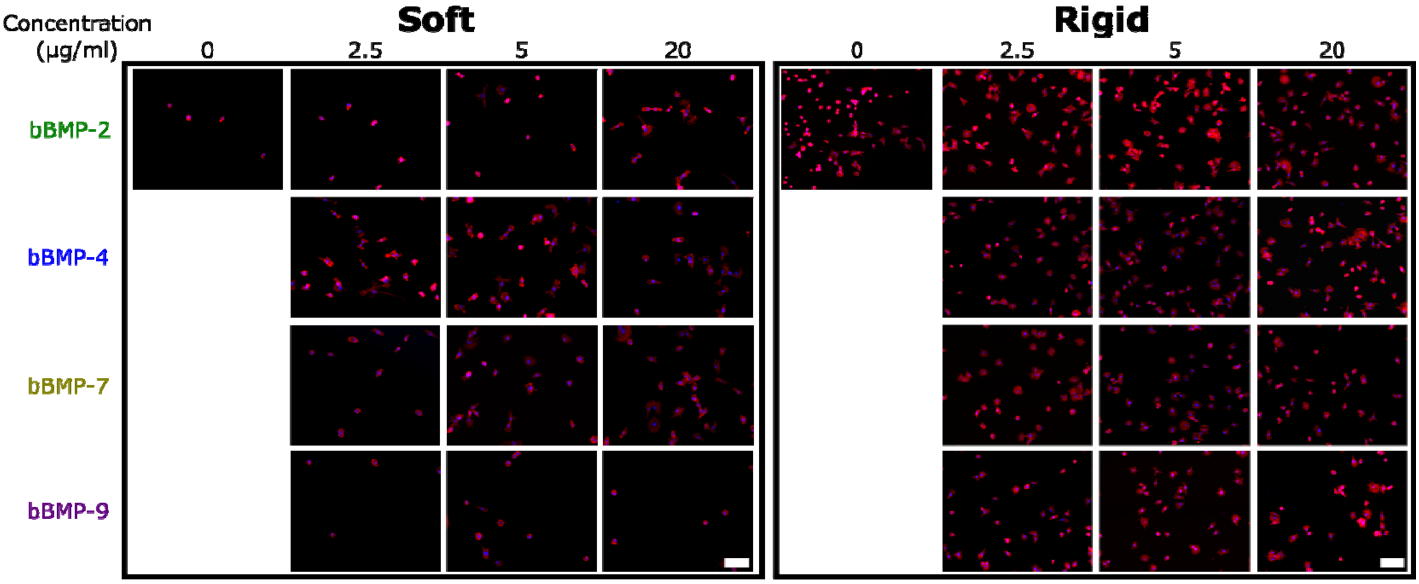

**Figure SI 2. Normalized cell number and cell area values to the non-BMP condition on C2C12 seeded on soft and rigid films with different BMP concentrations.** (**a**) Calculation of cell number per mm$^2$ of substrate area, and (**b**) cell area both of them normalized by the non-BMP condition. Thus, the effect of each bBMP for each film stiffness is exalted. Data represent three independent biological replicates with two wells per condition in each independent experiment (technical replicates). Statistical tests were done using non-parametric Kruskal-Wallis ANOVA test to compare soft (light color) and rigid (dark color) films (*$p < 0.05$; **$p < 0.01$).

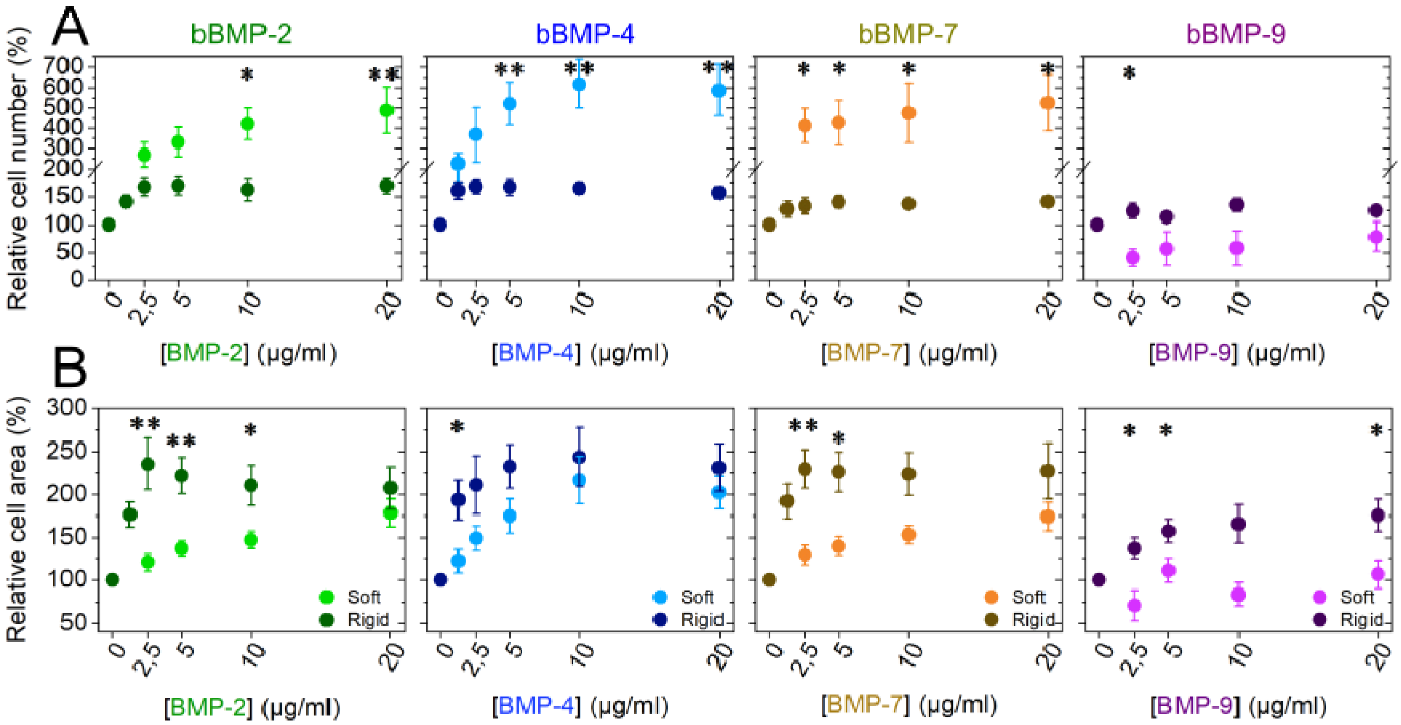

**Figure SI 3. Effects of soluble BMPs (sBMP-2, -4, -7, -9) on cell adhesion and spreading of C2C12 cells.** Cells were fixed after 5h, on soft films, and after 4h on rigid ones.. (**a**) The cell number (cells/mm$^2$ of substrate area) and (**b**) the cell spreading area (µm$^2$) are given. Data represent the mean ± SEM, with 3 independent biological replicates and 2 technical replicates in each experiment. Statistical tests were done using non-parametric Kruskal-Wallis ANOVA test to compare each BMP condition with the no BMP condition. * p ≤ 0.05.

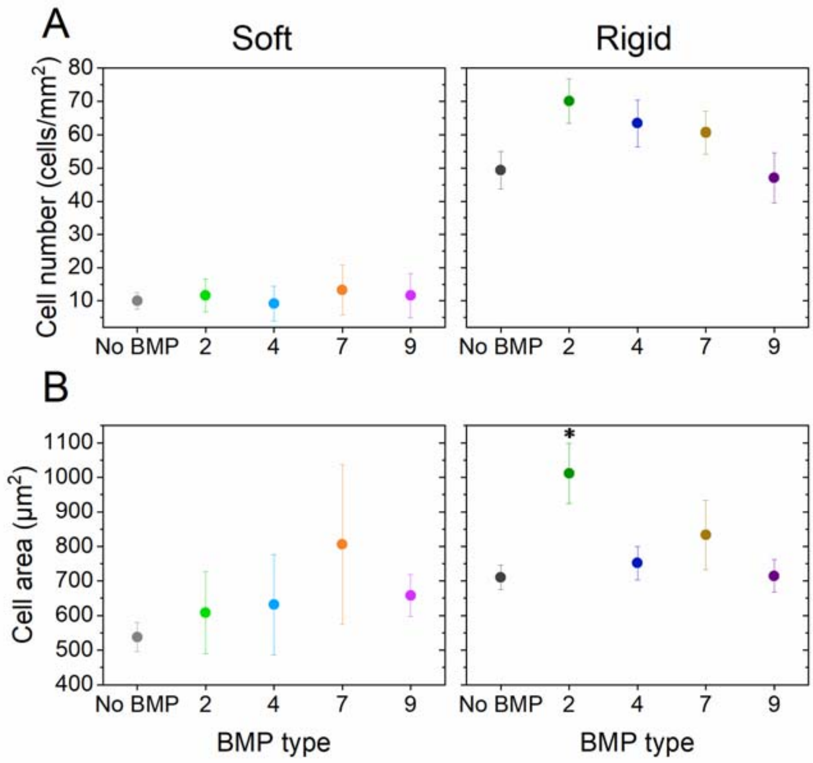

**Figure SI 4. Quantification of hPDSCs adhesion on soft and rigid films with four bBMPs (2, 4, 7, 9) at increasing BMP-loaded concentrations.** (**a**) Representative images of the cell actin cytoskeleton (red) on soft films 5h after cell seeding, and on rigid films 4h after cell seeding. (**b)** Cell number par mm$^2$ of substrate area and (**c**) cell area as a function of the BMP concentration in solution are shown. Quantitative parameters extracted from the fit of the experimental curves: (**d** and **e**) cell number and (**f** and **g**) cell area. Data represent the mean $\pm$ SEM of 3 independent experiments with 2 samples per condition in each experiments (at least a total of 400 cells analyzed per condition). Statistical tests were done between soft (light color) and rigid (dark color) films using non-parametric Kruskal-Wallis ANOVA test (*$p < 0.05$; **$p < 0.01$). Scale bar=200 µm.

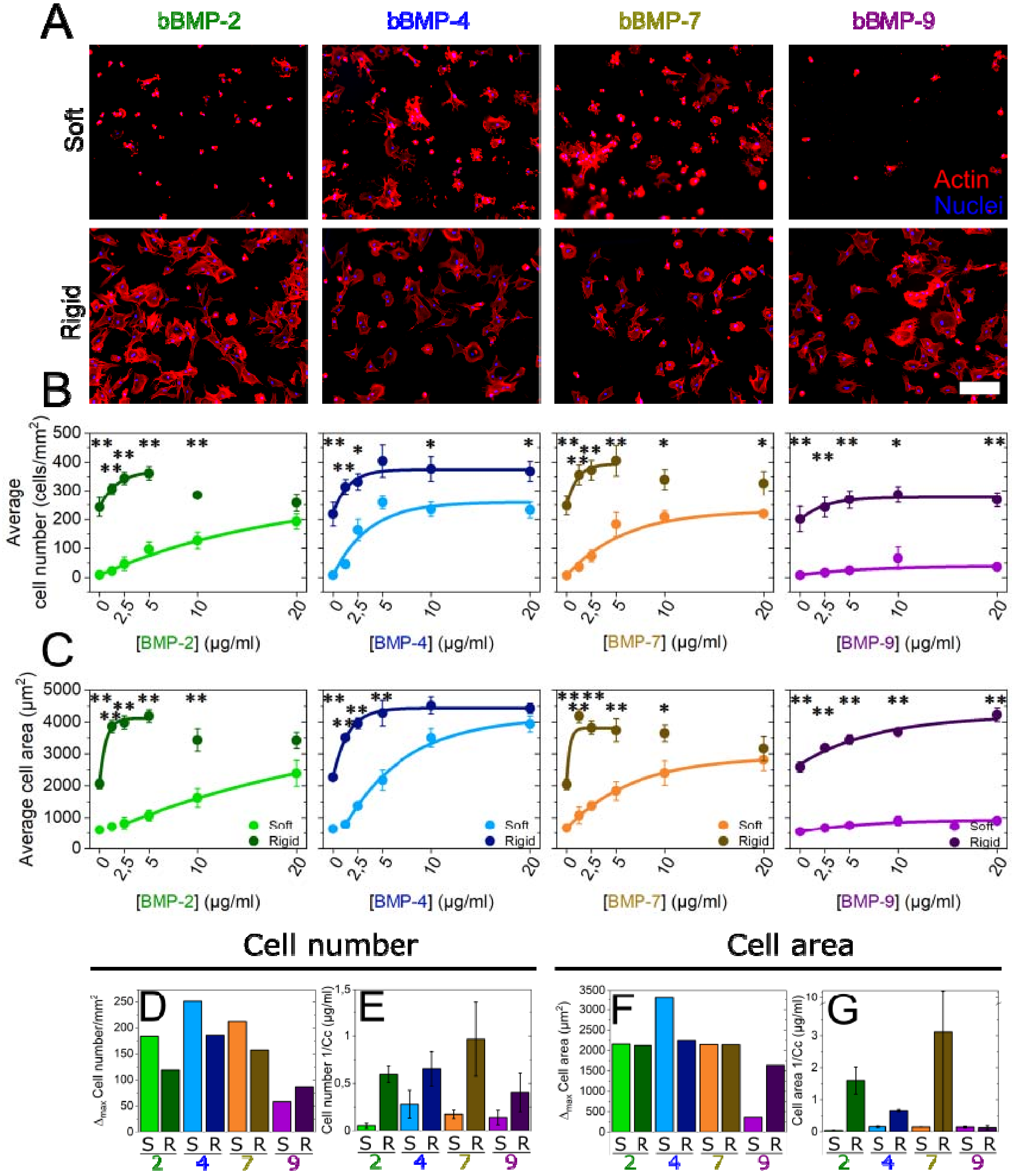

**Figure SI 5. Normalized cell number and cell area values to the non-BMP condition on hPDSCs seeded on soft and rigid films with different BMP concentrations.** (**a**) Calculation of cell number per mm$^2$ of substrate area and (**b**) cell area both of them normalized by the non-BMP condition. Thus, the effect of each bBMP for each film stiffness is exalted. Data represent three independent biological replicates with two wells per condition in each independent experiment (technical replicates). Statistical tests were done using non-parametric Kruskal-Wallis ANOVA test to compare soft (light color) and rigid (dark color) films (*p < 0.05; **p < 0.01).

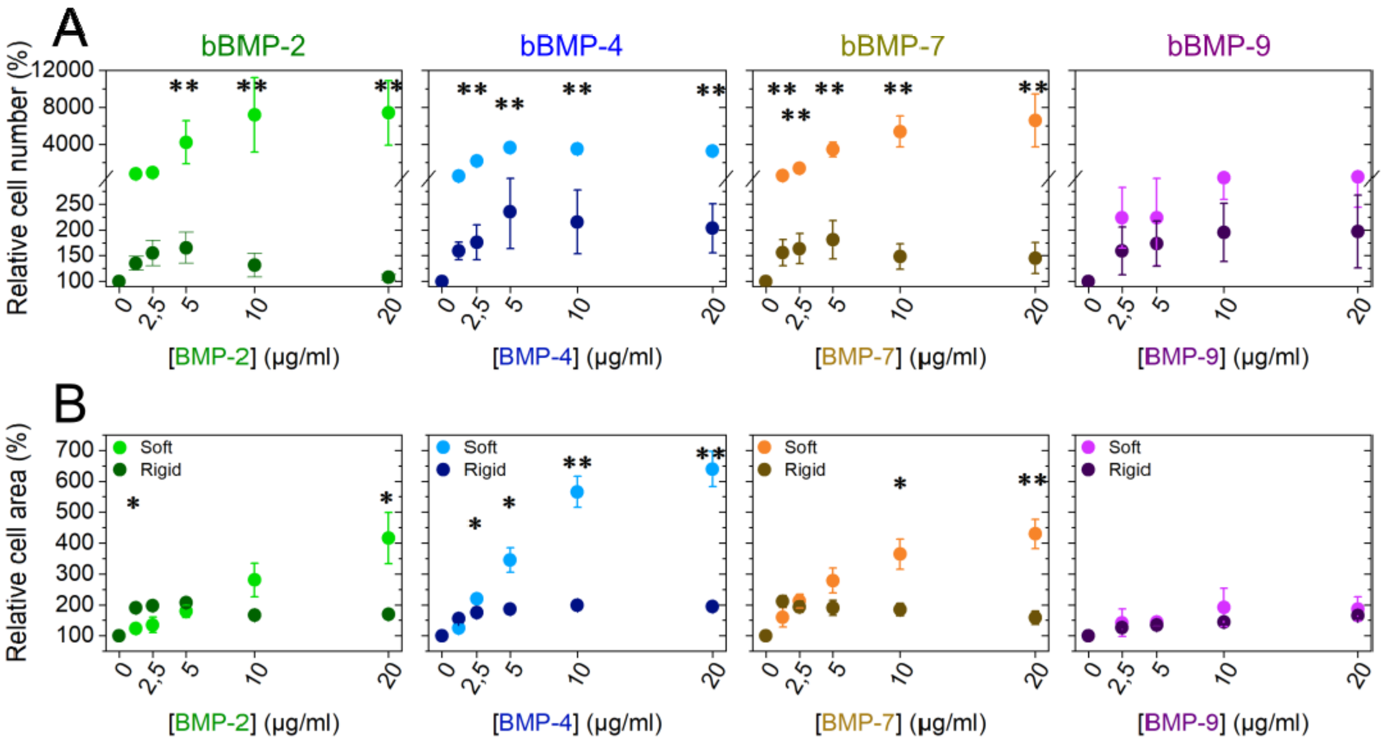

**Figure SI 6. Effects of soluble BMPs (sBMP-2, -4, -7, -9) on cell adhesion and spreading of hPDSCs cells.** Cells were fixed after 5h, on soft films, and after 4h on rigid ones. (**a**) The cell number (cells/mm$^2$ of substrate area) and (**b**) the cell spreading area (µm$^2$) are given. Data represent the mean ± SEM, with 3 independent biological replicates and 2 technical replicates in each experiment. Statistical tests were done using non-parametric Kruskal-Wallis ANOVA test to compare each BMP condition with the non BMP condition. * $p \leq 0.05$.

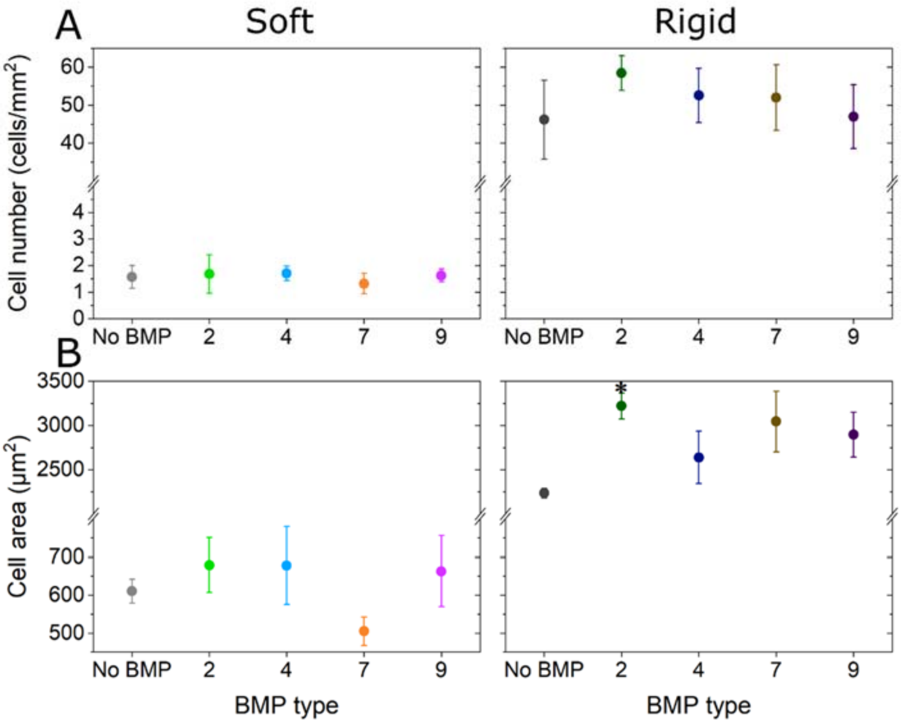

**Figure SI7. Representative example of high and low pSMAD fluorescent signal.** Boxplot showing an example of pSMAD1,5,9 signal distribution of individual cells, represented by the small points. No BMP condition vs bBMP-9 at 20µg/ml on rigid films are represented. Representative fluorescent images of each condition, representing a cell with low pSMAD1,5,9 signal, seeded on no-BMP condition, and a cell with a high pSMAD1,5,9 signal, seeded on bBMP-9 at 20µg/ml. Both conditions correspond to a rigid film. Immunofluorescent staining was performed 4h after cell seeding. Box plots represent the 25 and 75% percentiles. The horizontal line in the box represent the median (50% percentile). The filled square represent the mean value, and the error bars represent the standard deviation. Scale bar=25 µm.

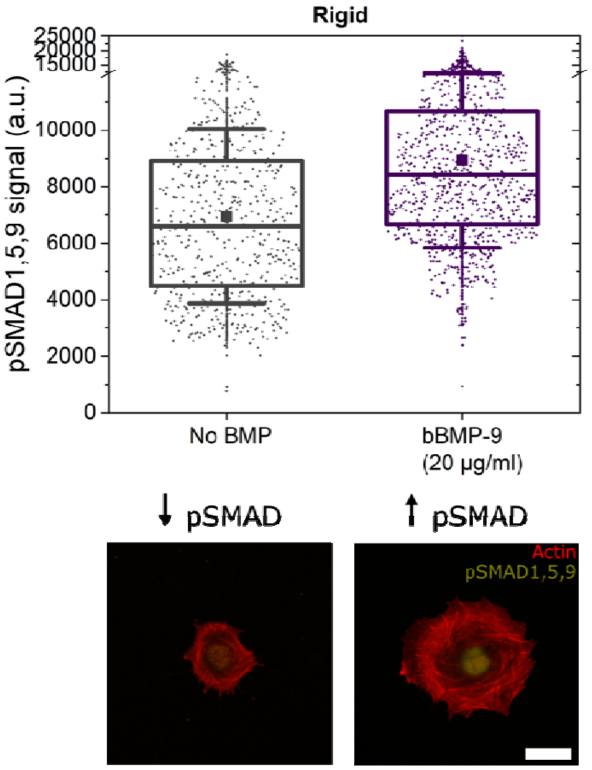

**Figure SI8. pSMAD analyses for the four bBMPs (2, 4, 7 and 9) on soft and rigid films.** Fluorescent signal quantification of (**a**) pSMAD1,5,9 and (**b**) pSMAD2 as a function of the BMP concentration in soft (light color) and rigid films (dark color). Data represent the mean ± SEM of 2 to 4 independent experiments with two samples per condition in each independent experiment (at least 400 cells were analyzed in total). Statistical tests were done using non-parametric Kruskal-Wallis ANOVA test to compare soft and rigid films. However, no statistical differences were found.

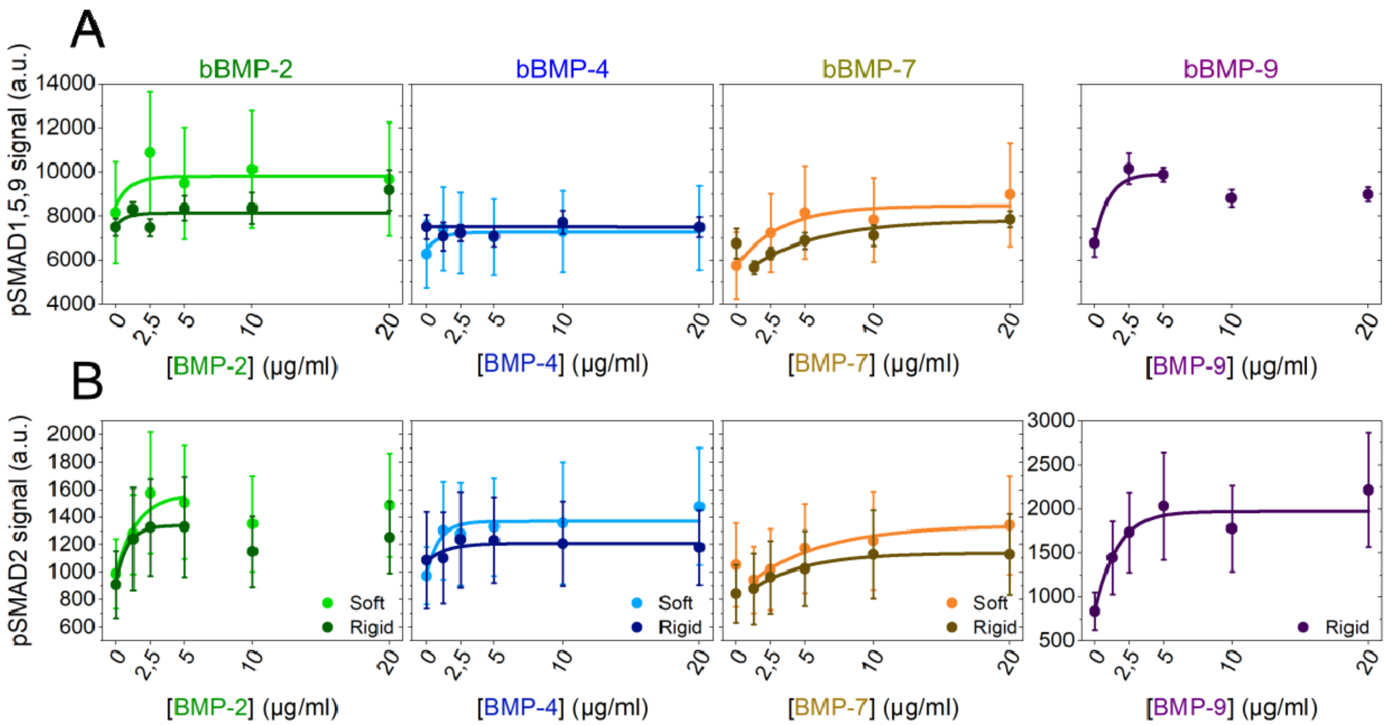

**Figure SI 9. Effects of soluble BMPs (sBMP-2, -4, -7, -9) on pSMAD1,5,9 signal on C2C12 cells.** Cells were fixed after 5h, on soft films, and after 4h on rigid ones. Data represent the mean ± SEM, with 3 independent biological replicates and 2 technical replicates in each experiment. Statistical tests were done using non-parametric Kruskal-Wallis ANOVA test to compare each BMP condition with the no-BMP condition. * $p \leq 0.05$.

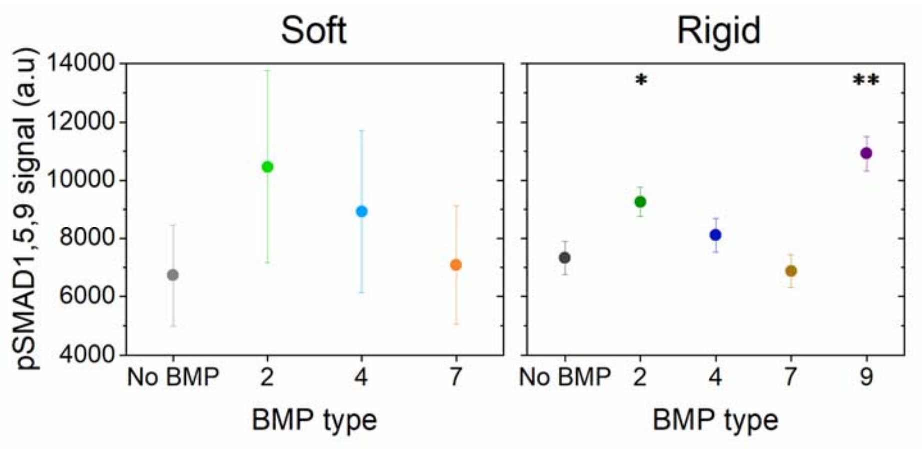

**Figure SI 10**. **Representative images of an ALP staining for C2C12 cells on soft and rigid films with all 4 bBMPs (2, 4, 7, 9).** C2C12 cells were fixed and stained 3 days after cell seeding to measure ALP activity. The loading BMP concentration of 20 µg/ml for all BMPs, and the non-BMP condition are shown as representative examples.

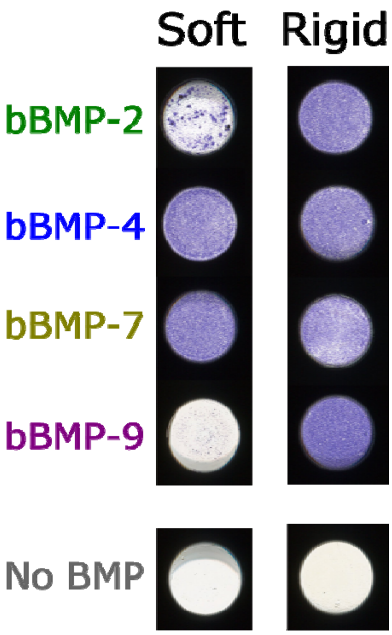

**FIGURE SI 11. Identification of C2C12 cells-specific BMP receptors and integrins, using the ENCODE database.** Pie chart of the percentage of expression of BMP receptor type I, BMP receptor type II and β integrins (1, 3 and 5), illustrating the predominance of certain adhesion receptors in C2C12 cells. Data were obtained by analyzing RNA sequencing data made for the ENCODE public research project.

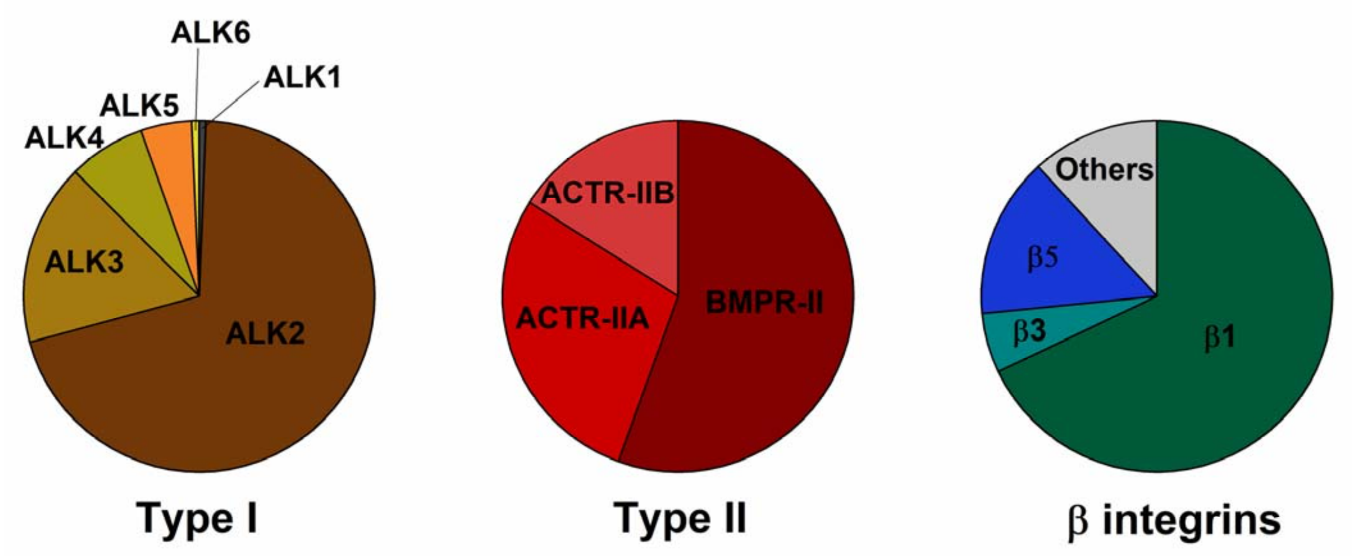

**Figure SI 12**. **C2C12 transfection control.** The gene expression level of each knocked down gene, as well as the corresponding scrambled condition, are shown for the BMP receptors type I, type II and the three β integrins studied. All the receptors were correctly silenced, with expression levels below 10% with respect to the scrambled condition. Bars represent the mean and error bars represent the ± SEM, with 3 biological replicates and 2 technical replicates per experiment.

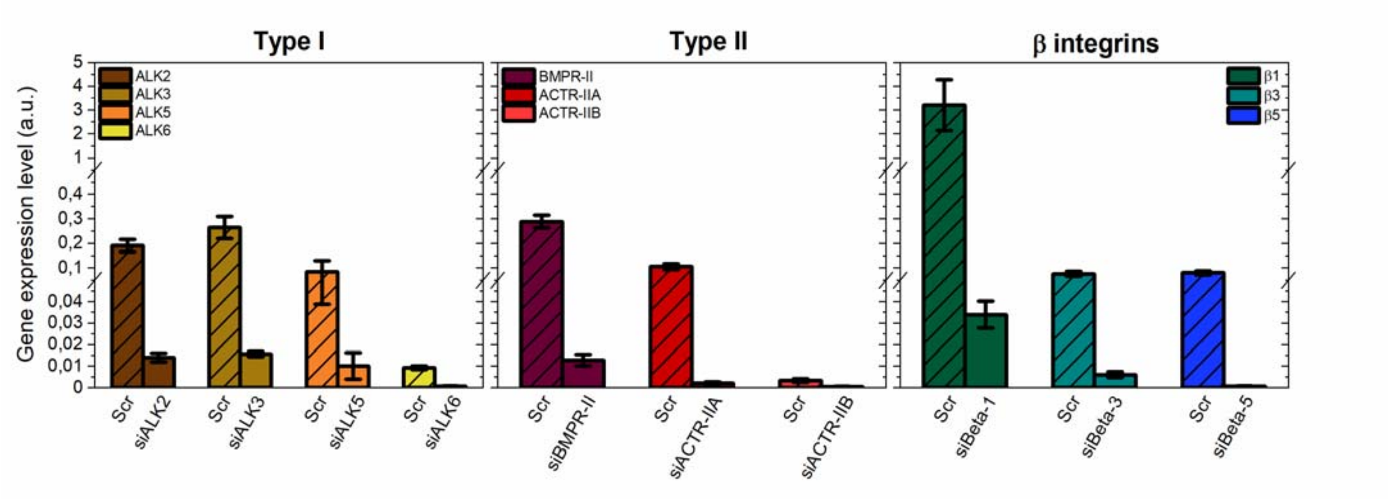

**Figure SI 13. Effect of BMP receptor and beta chain integrin silencing on cell adhesion and spreading on rigid films with no BMPs.** C2C12 cells were transfected with siRNA against BMP receptor type I (ALK2, ALK3, ALK5, ALK6), BMP receptor type II (BMPR-II, ACTR-IIA, ACTR-IIB) and beta chain integrins (β1, β3, β5) and were plated on rigid films without bBMPs for 4 h. The cell number par mm$^2$ of substrate area and the spreading area were quantified, and the relative % is given, in comparison to a control scrambled siRNA. (**a**) Relative cell number (%), (**b**) relative cell area (%). Data represent the mean ± SEM, with 2 to 4 biological replicates and 2 technical replicates per experiment.

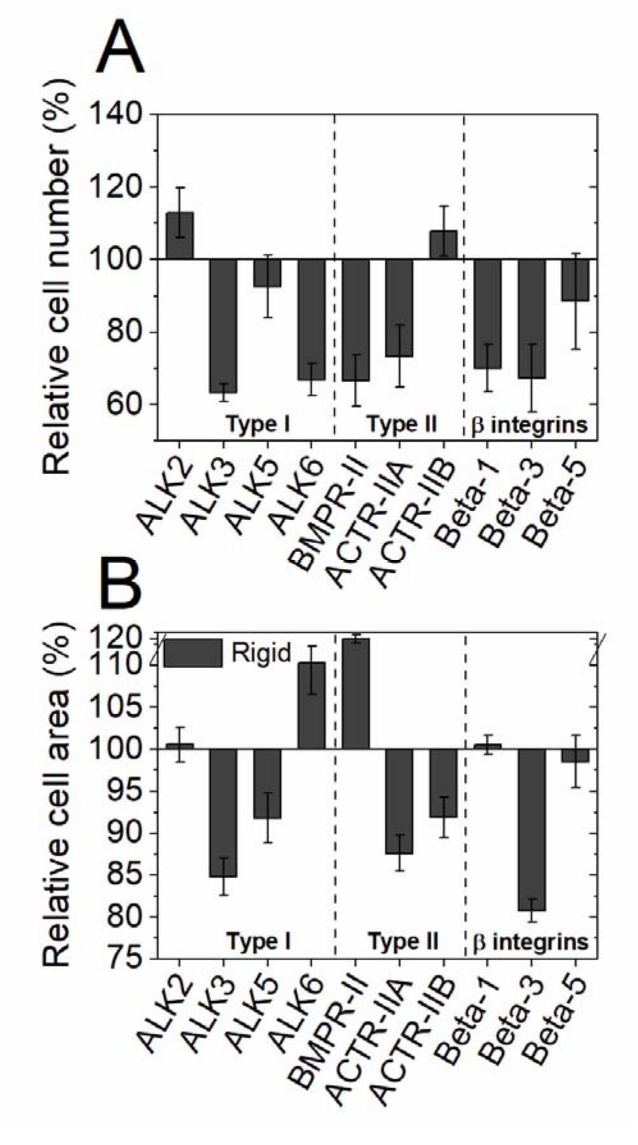

**Figure SI 14. Non normalized data showing the effect of BMP receptor and beta chain integrin silencing on cell adhesion and spreading.** C2C12 cells were transfected with siRNA against BMP receptor type I (ALK2, ALK3, ALK5, ALK6), BMP receptor type II (BMPR-II, ACTR-IIA, ACTR-IIB) and beta chain integrins (β1, β3, β5) and were plated on soft and rigid films with or without bBMPs for 5 or 4 h, respectively. (**a**) The cell number par mm$^2$ of substrate area and (**b**) the spreading area were quantified. Data represent the mean ± SEM, with 2 to 4 biological replicates and 2 technical replicates per experiment. Statistical tests were done using the non-parametric Kruskal-Wallis ANOVA test (*, $p \leq 0.05$; **, $p \leq 0.01$). Statistical comparisons were made between the different knock down conditions and the scrambled.

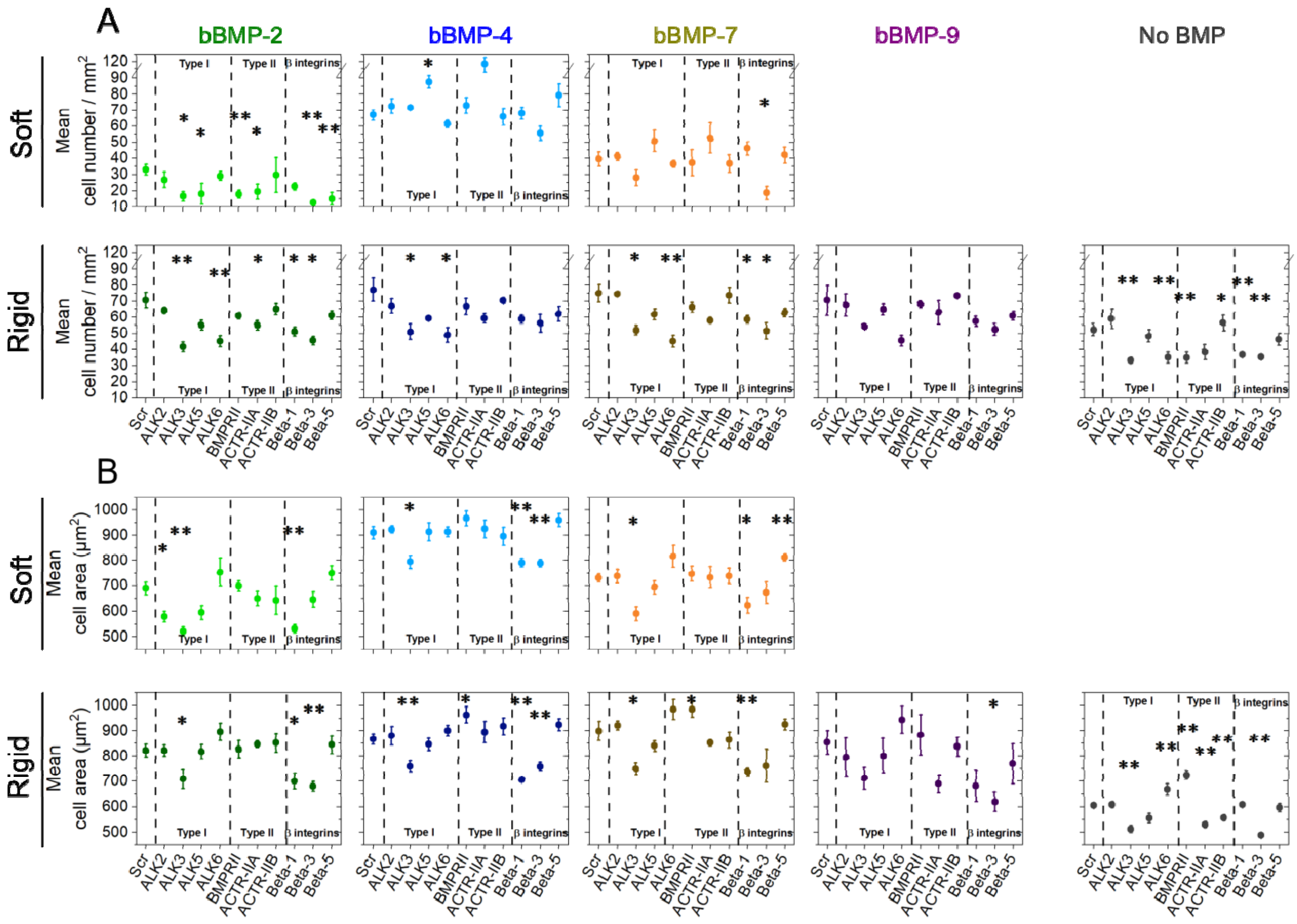

**Figure SI 15. Effect of BMP receptor and beta chain integrin silencing on cell differentiation to bone, on rigid films with no BMPs.** C2C12 cells were transfected with siRNA against BMP receptor type I (ALK2, ALK3, ALK5, ALK6), BMP receptor type II (BMPR-II, ACTR-IIA, ACTR-IIB) and beta chain integrins (β1, β3, β5) and were plated on rigid films without bBMPs for 4 h. (**a**) pSMAD1,5,9 and (**b**) pSMAD2 signal, as well as (**c**) ALP activity were quantified. The relative % is given, in comparison to a control scrambled siRNA. Data represent the mean ± SEM, with 2 to 4 biological replicates and 2 technical replicates per experiment.

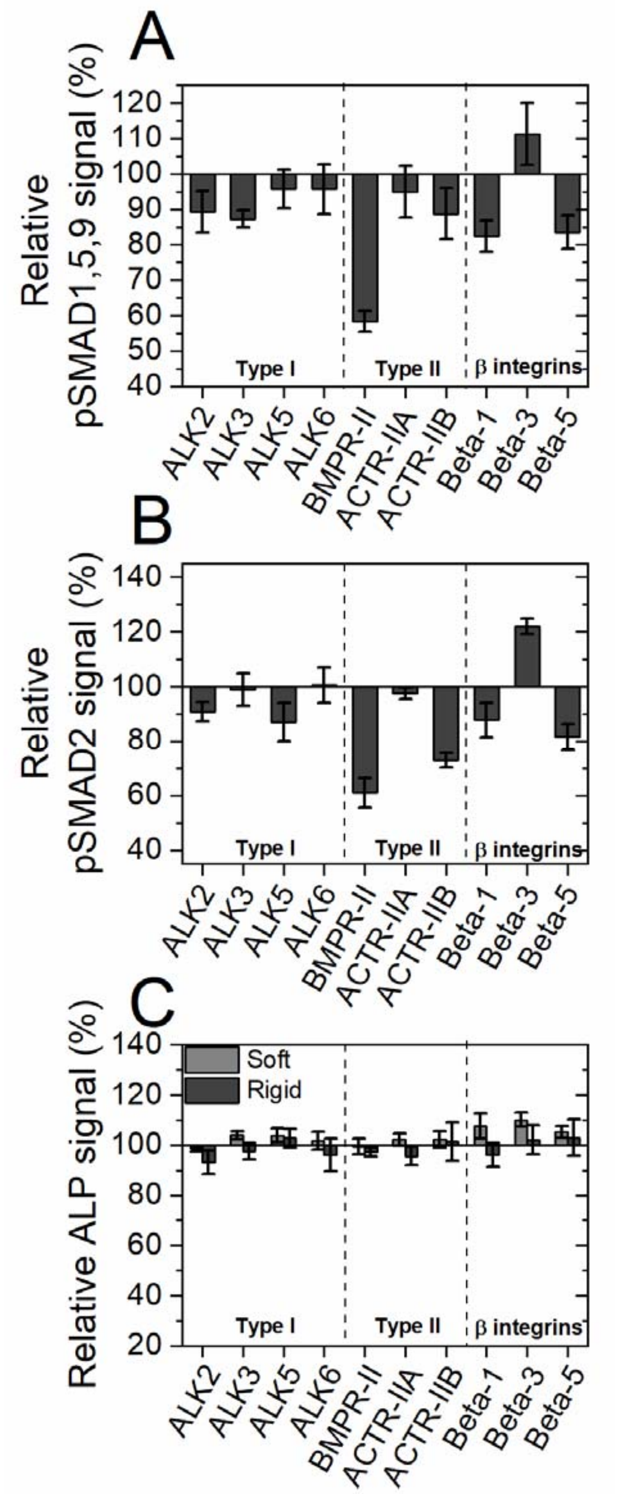

**Figure SI 16**. **Non normalized data showing the effect of BMP receptor and beta chain integrin silencing on pSMAD1,5,9, pSMAD2 and ALP activation.** C2C12 cells were transfected with siRNA against BMP receptor type I (ALK2, ALK3, ALK5, ALK6), BMP receptor type II (BMPR-II, ACTR-IIA, ACTR-IIB) and beta chain integrins (β1, β3, β5) and were plated on soft and rigid films with or without bBMPs for 5 or 4 h, respectively. (**a**) The pSMAD1,5,9 and (**b**) pSMAD2 fluorescent signals, together with (**c**) the ALP activation level were quantified. Data represent the mean ± SEM, with 2 to 4 biological replicates and 2 technical replicates per experiment. Statistical tests were done using non-parametric Kruskal-Wallis ANOVA test (*, p ≤ 0.05; **, p ≤ 0.01). Statistical comparisons were made between the different knock down conditions and the scrambled condition.

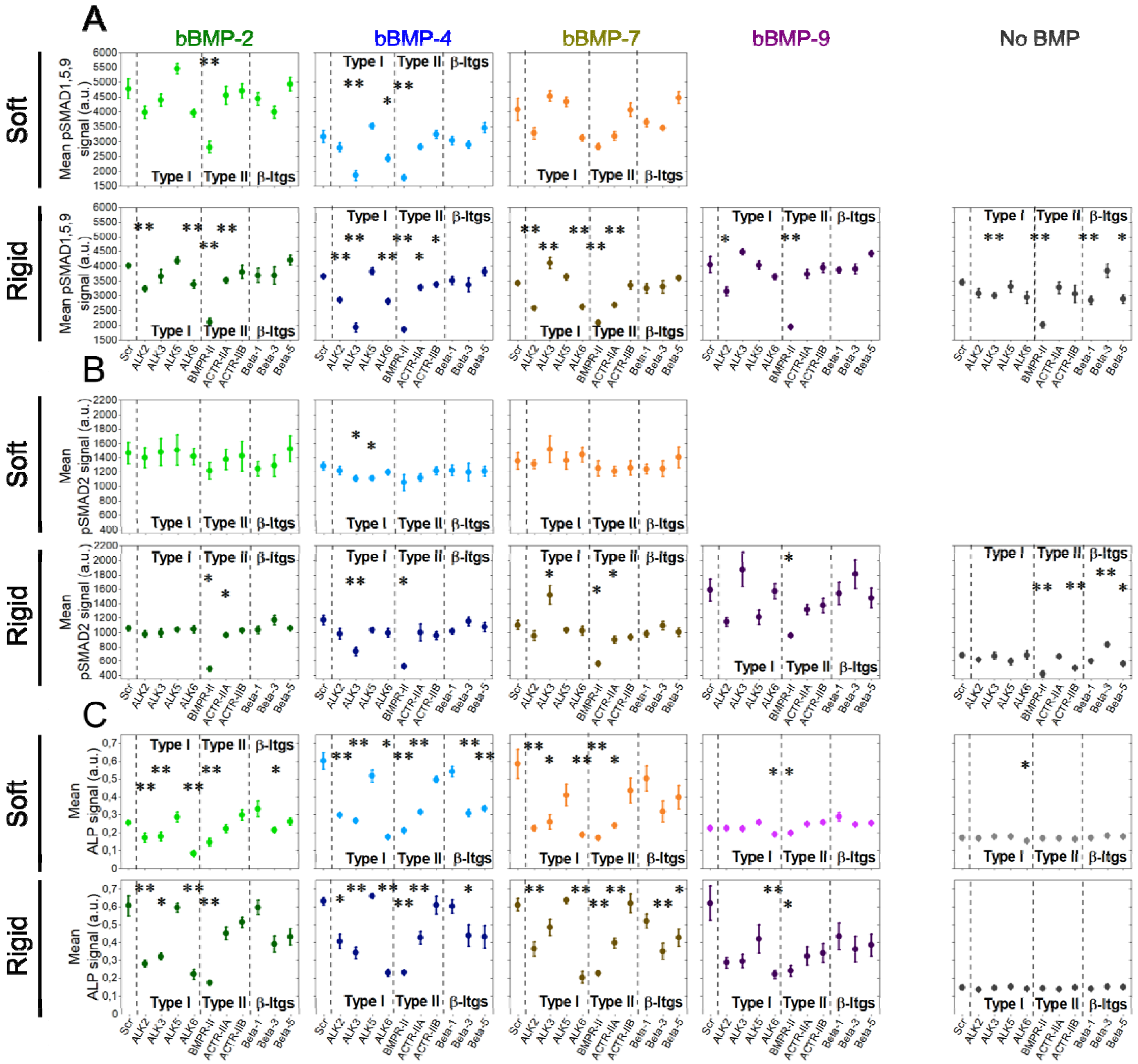

**Figure SI 17**. **Representative pictures of an ALP staining on C2C12 cells for each bBMP (2, 4, 7 and 9) and each knock down condition, on soft and rigid films.** C2C12 cells were fixed 3 days after seeding, and were stained for ALP, on soft and rigid films with a BMP loading concentration of 20 µg/ml, as well as on the non-BMP control condition.

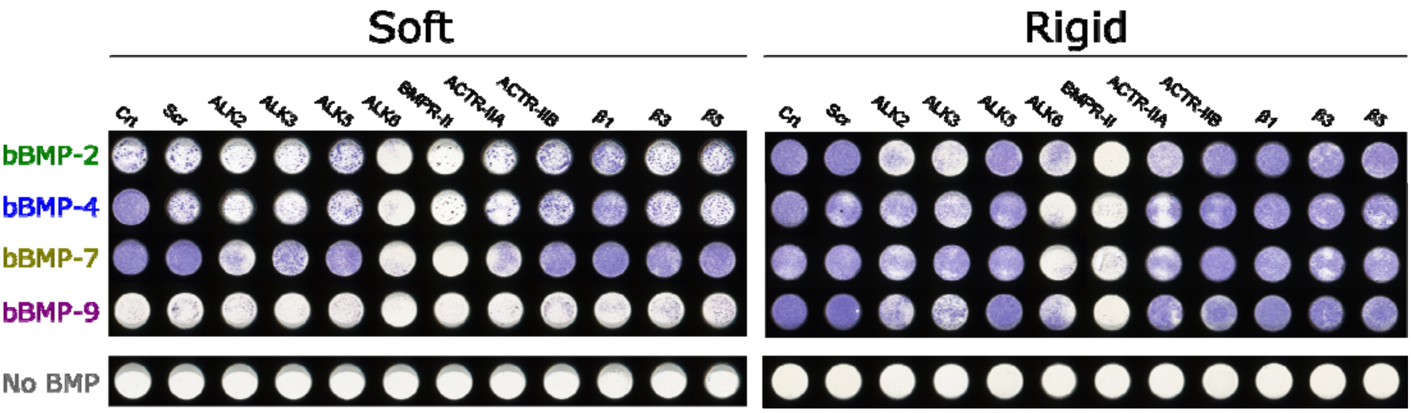
